# Identification of functionally-distinct macrophage subpopulations regulated by efferocytosis in *Drosophila*

**DOI:** 10.1101/2020.04.17.047472

**Authors:** Jonathon Alexis Coates, Amy Brittle, Emma Louise Armitage, Martin Peter Zeidler, Iwan Robert Evans

## Abstract

Macrophages are a highly heterogeneous population of cells, with this diversity stemming in part from the existence of tissue resident populations and an ability to adopt a variety of activation states in response to stimuli. *Drosophila* blood cells (hemocytes) are dominated by a lineage of cells considered to be the functional equivalents of mammalian macrophages (plasmatocytes). Until very recently plasmatocytes were thought to be a homogeneous population. Here, we identify enhancer elements that label subpopulations of plasmatocytes, which vary in abundance across the lifecourse of the fly. We demonstrate that these plasmatocyte subpopulations behave in a functionally-distinct manner when compared to the overall population, including more potent migratory responses to injury and decreased clearance of apoptotic cells within the developing embryo. Additionally, these subpopulations display differential localisation and dynamics in pupae and adults, hinting at the presence of tissue-resident macrophages in the fly. Our enhancer analysis also allows us to identify novel candidate genes involved in plasmatocyte behaviour in vivo. Misexpression of one such enhancer-linked gene (*calnexin14D*) in all plasmatocytes improves wound responses, causing the overall population to behave more like the subpopulation marked by the *calnexin14D*-associated enhancer. Finally, we show that, we are able to modulate the number of cells within some subpopulations via exposure to increased levels of apoptotic cell death, thereby decreasing the number of plasmatocytes within more wound-responsive subpopulations. Taken together our data demonstrates the existence of macrophage heterogeneity in *Drosophila* and identifies mechanisms involved in the specification and function of these plasmatocyte subpopulations. Furthermore, this work identifies key molecular tools with which *Drosophila* can be used as a highly genetically-tractable, in vivo system to study the biology of macrophage heterogeneity.

## Introduction

Macrophages are key innate immune cells responsible for clearing infections, debris and apoptotic cells, the promotion of wound healing and are necessary for normal development [1]. However, their aberrant behaviour can also cause or exacerbate numerous human disease states, including cancer, atherosclerosis and neurodegeneration [1]. Macrophages are a highly heterogeneous population of cells, which enables them to carry out their wide variety of roles, and this heterogeneity arises from diverse processes. These processes include the dissemination and maintenance of tissue resident populations [2] and the ability to adopt a spectrum of different activation states (termed macrophage polarisation), which can range from pro-inflammatory (historically termed as M1-like) to anti-inflammatory, pro-healing (M2-like) macrophage activation states [3,4].

Macrophage heterogeneity appears to be conserved across jawed vertebrate lineages. Evidence suggests the existence of pro-inflammatory macrophage populations [5] and myeloid-derived microglia in zebrafish [6,7], with polarisation also a well-defined phenomenon in other fish species [8]. Vertebrate macrophages interact with and can become polarised in response to signals produced by Th1 and Th2 cells, leading to acquisition of M1-like and M2-like activation states, respectively. To date this form of heterogeneity has been considered to be restricted to organisms containing both an adaptive and an innate immune system. B and T cell-based adaptive immunity is thought to have evolved in teleost fish [9] and the diversity of macrophage populations in organisms possessing only an innate immune system appears more restricted. However, even comparing mammals as closely related as mice and humans, macrophage markers can be highly divergent [10], therefore other approaches and markers might be required to identify equivalent macrophage diversity in lower organisms.

Macrophage heterogeneity has been extensively studied in mammalian systems and, although this has provided a good understanding of how macrophages determine their polarisation state, this has also identified considerable complexity with many activation states possible [11]. Additional complexity arises with both M1-like and M2-like macrophages found at the same sites of pathology, for example within atherosclerotic plaques [12]. Furthermore, the cytokine profiles that can be induced in vitro depend on the exact activation methods used experimentally and these do not necessarily reflect polarisation states in vivo [13], while other macrophage subpopulations may be missed by in vitro approaches. Given these intricacies, it is clear that we still need to better understand the fundamental components and pathways responsible for the specification of different macrophage subtypes, particularly in vivo. Recently the “macrophage-first” hypothesis has been proposed, re-emphasising the idea that acute signals polarise macrophages ahead of the involvement of T cells [8]. Consequently, organisms without a fully-developed adaptive immune system represent intriguing models in which to examine this idea and better understand macrophage heterogeneity in vivo.

*Drosophila melanogaster* has been extensively used to study innate immunity [14], but lacks an adaptive immune system. Fruit flies possess three types of blood cell (also referred to as hemocytes): plasmatocytes, crystal cells and lamellocytes. Of these, plasmatocytes are functionally equivalent to vertebrate macrophages [15,16], with the capacity to phagocytose apoptotic cells and pathogens, secrete extracellular matrix, disperse during development and migrate to sites of injury [17]. Although *Drosophila* blood lineages are considerably less complex than their vertebrate equivalents, they are specified via transcription factors related to those used during vertebrate myelopoiesis, including GATA and Runx-related proteins [15]. Furthermore, plasmatocytes utilise evolutionarily-conserved genes in common with vertebrate innate immune cells to migrate (e.g. SCAR/WAVE, integrins and Rho GTPases [18–22]) and phagocytose (e.g. the CED-1 family member Draper [23] and CD36-related receptor Croquemort [24]). Given these striking levels of functional and molecular conservation, *Drosophila* has been extensively used for research into macrophage behaviour in vivo with its genetic tractability and in vivo imaging capabilities facilitating elucidation of different macrophage behaviours conserved through evolution [16,17]. However, despite these evolutionarily-conserved commonalities, the plasmatocyte lineage has, until very recently, been considered a homogeneous cell population. Hints that *Drosophila* plasmatocytes may exhibit heterogeneity exist in the literature with variation in marker expression observed in larval hemocytes [25] and non-uniform expression of TGF-β homologues upon injury or infection in adults [26]. Recent single-cell RNA-sequencing (scRNA-seq) experiments performed on larval hemocytes demonstrated the presence of multiple clusters of cells, which were interpreted as representing either different stages of differentiation or functional groupings [27,28]. However, the in vivo identification of subtypes and insights into the roles and specification mechanisms of potential macrophage subtypes in *Drosophila* has not yet been described.

Here, we describe the first identification and characterisation of molecularly and functionally-distinct plasmatocyte subpopulations within *Drosophila melanogaster*. Drawing on a collection of reporter lines [29], we have identified regulatory elements that define novel plasmatocyte subpopulations in vivo. We show that these molecularly-distinct subpopulations exhibit functional differences compared to the overall plasmatocyte population and that the proportions of cells within these subpopulations can be modulated by external stimuli such as increased levels of apoptosis. Furthermore, we show that misexpression of a gene associated with a subpopulation-specific enhancer element is able to modulate plasmatocyte behaviour in vivo, thereby identifying novel effector genes of plasmatocyte subpopulation function. Together our findings reveal that macrophage heterogeneity is a fundamental and evolutionarily-conserved characteristic of innate immunity that pre-dates the development of the adaptive immune system. This significantly extends the utility of an already powerful genetic model system and provides further avenues to understand regulation of innate immunity and macrophage heterogeneity.

## Results

### *Drosophila* embryonic plasmatocytes do not behave as a uniform population of cells

The macrophage lineage of hemocytes (plasmatocytes) has historically been considered a homogeneous population of cells. However, careful analysis of plasmatocyte behaviour in vivo suggested to us that this lineage might not be functionally uniform. For instance, imaging the inflammatory responses of plasmatocytes to epithelial wounds, we find that some cells close to injury sites rapidly respond by migrating to the wound, while other neighbouring cells fail to respond, (Figure 1a; Supplementary Movie 1). We also find that plasmatocytes exhibit variation in their expression of well-characterised plasmatocyte markers (*crq-GAL4* [19,24]; Figure 1b-b’) and display a broad diversity in their migration speeds within the embryo (random migration at stage 15, Figure 1c-d). These professional phagocytes also display differences in their capacities to phagocytose apoptotic cells with some cells engulfing many apoptotic particles, whereas others engulf very few, if any (Figure 1e). Furthermore, phagocytosis of microorganisms by larval hemocytes also varies significantly from cell-to-cell in vitro (Figure 1f). These differences within the plasmatocyte lineage led us to hypothesise that this cell population is more heterogeneous than previously appreciated.

**Figure 1.**
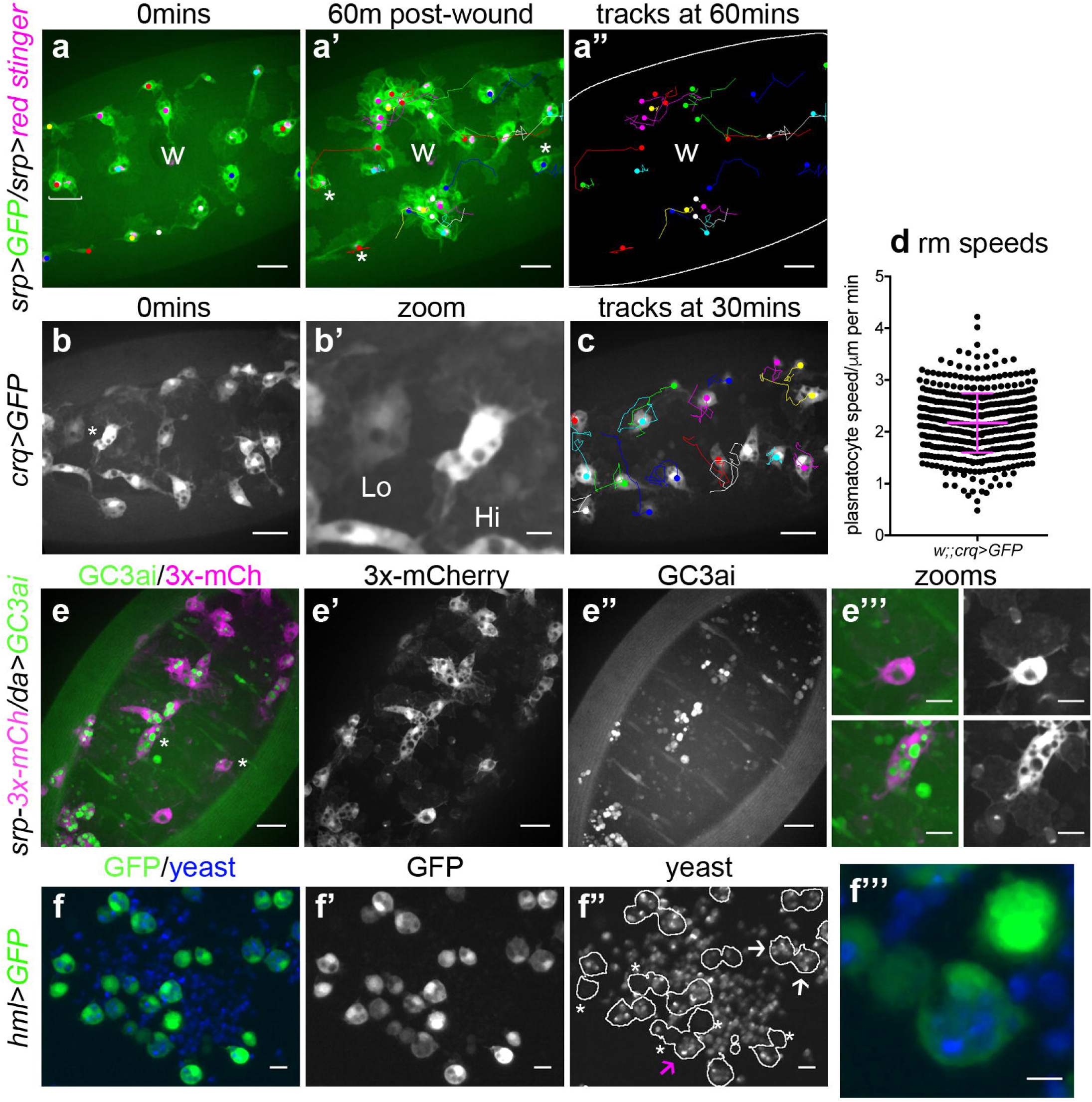
Heterogeneity of *Drosophila* embryonic plasmatocyte responses. (a) GFP (green) and nuclear red stinger (magenta) labelled plasmatocytes on the ventral side of a stage 15 embryo at 0-minutes (a) and 60-minutes post-wounding (a’); plasmatocyte tracks at each timepoint are overlaid and shown in full in a’’. Examples of plasmatocytes that fail to respond to the wound indicated via asterisks; “w” shows centre of the wound; square bracket in (a) shows neighbouring plasmatocytes, one of which responds to wounding, the other fails to respond (see Supplementary Movie 1). (b) imaging of plasmatocytes labelled using *crq-GAL4* to drive expression of GFP reveals a wide range in levels of *crq* promoter activity within plasmatocytes at stage 15; (b’) shows zoom of cells marked by an asterisk in (b). (c) overlay of plasmatocyte tracks of cells shown in (b) showing significant variation in their random migration speeds. (d) scatterplot of plasmatocyte random migration (rm) speeds (taken from 23 embryos); line and error bars show mean and standard deviation, respectively. (e) imaging the ventral middle at stage 15 shows a wide range in the amount of apoptotic cell clearance (green in merge, labelled via the caspase-sensitive reporter GC3ai) undertaken by plasmatocytes (magenta in merge, labelled via *srp-3x-mCherry* reporter); (e’) and (e’’) show mCherry and GC3ai channels alone; (e’’’) shows zoomed examples of cells devoid/full of engulfed GC3ai particles, which are indicated by asterisks in (e). (f) larval hemocytes (green in merge, labelled via *hml(Δ)-GAL4* driven expression of GFP) exhibit a range in their capacities to engulf calcofluor-labelled yeast (blue in merge) in vitro; (f’) and (f’’) show GFP and yeast channels alone, respectively; white lines indicate cell edges in (f’’); asterisks in (f’’) indicate cells that have failed to phagocytose yeast; white arrows in (f’’) indicate cells that have phagocytosed multiple yeast particles; magenta arrow in (f’’) indicates zoomed region shown in (f’’’). Scale bars represent 20μm (a-a’’, b, c, e-e’’), 10μm (e’’’, f-f’’), or 5μm (b’, f’’’). See Supplementary Table 1 for full list of genotypes.

### Discrete subpopulations of plasmatocytes are present in the developing *Drosophila* embryo

Given the diversity in plasmatocyte behaviour (Figure 1), we hypothesised that macrophage heterogeneity represents an evolutionarily-conserved feature of innate immunity, which therefore originally evolved in the absence of an adaptive immune system. To address this and look for molecular differences between plasmatocytes, we examined transgenic enhancer reporter lines (*VT-GAL4* lines) produced as part of a recent large-scale tilling array screen [29] that had been annotated as labelling hemocytes (http://enhancers.starklab.org/). Based on examination of the published *VT-GAL4* expression patterns, we identified *VT-GAL4* lines that appeared to label reduced numbers of plasmatocytes in the embryo, reasoning that plasmatocyte subpopulations could be molecularly identified on the basis of differences in reporter expression. While a number of the enhancers appeared to label all plasmatocytes (e.g. *VT41692-GAL4*), we identified several that labelled discrete numbers of plasmatocytes (Figure 2a). We next confirmed that the cells labelled by these *VT-GAL4* lines were plasmatocytes by using these constructs to drive expression of *UAS-tdTomato* in the background of a *GAL4*-independent, pan-hemocyte marker (*srp-GMA* (GFP-tagged actin-binding domain of moesin); Figure 2b-d; [30]). As initially predicted based on their morphology and position during embryogenesis, each of the *VT-GAL4* lines marking potential subpopulations did indeed express in the hemocyte lineage Figure 2e). These subpopulation cells were identified as plasmatocytes based upon their morphology, the absence of lamellocytes in embryos and the non-migratory nature of crystal cells (Figure 2e; [16]) and could be observed to follow both the dorsal and ventral migration routes [17] used by these cells during their developmental dispersal (Figure 2e). In order to quantify the proportion of cells labelled by each *VT-GAL4* line, we counted the number of cells labelled on the ventral midline of the developing stage 15 embryo, using *VT-GAL4* lines to drive GFP expression. This verified reproducible and consistent labelling of discrete subsets of plasmatocytes (Figure 2f-h), suggesting that these cells represent stable subpopulations within this macrophage lineage.

**Figure 2.**
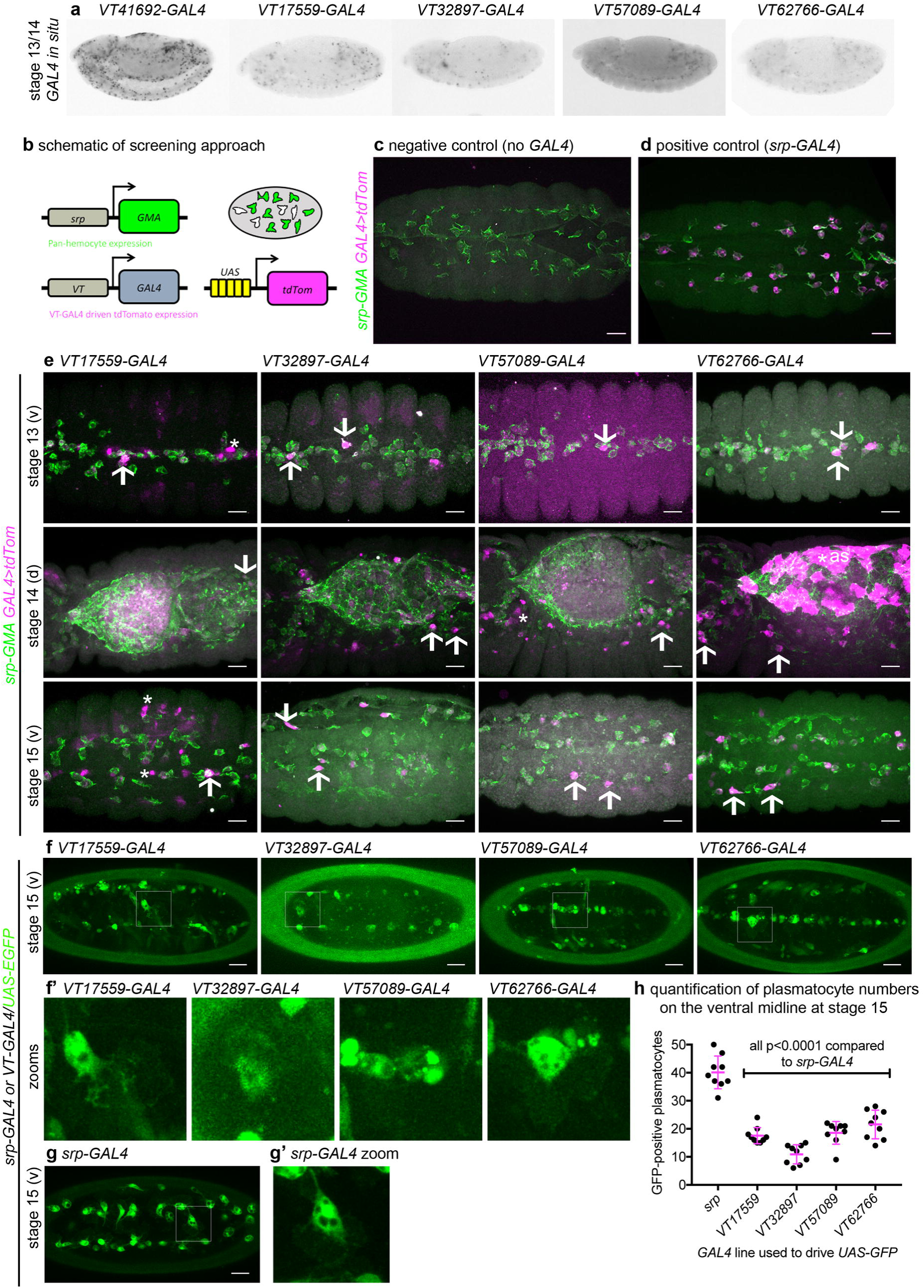
Enhancers labelling plasmatocyte subpopulations in *Drosophila*. (a) lateral views of stage 13/14 embryos with in situ hybridisation performed for *GAL4* for indicated *VT-GAL4* lines (anterior is left). Taken with permission from http://enhancers.starklab.org/ (n.b. Stark Lab retain copyright of these images); *VT41692-GAL4* represents an example in which the majority of plasmatocytes are labelled. (b) schematic diagram showing screening approach to identify subpopulations of plasmatocytes: *VT-GAL4* positive plasmatocytes will express both GMA (green) and tdTomato (magenta) – white cells in the schematic. (c-d) images showing the ventral midline at stage 14 of negative control (no driver; *w;UAS-tdTom/+;srp-GMA*) and positive control (w;*srp-GAL4/UAS-tdTom;srp-GMA*) embryos. (e) images showing embryos containing *VT-GAL4* labelled cells (via *UAS-tdTomato*, shown in magenta) at stage 13 (first row, ventral views), stage 14 (second row, dorsal views) and stage 15 (third row, ventral views). The entire hemocyte population is labelled via *srp-GMA* (green); arrows indicate examples of *VT-GAL4* positive plasmatocytes; asterisks indicate *VT-GAL4* positive cells that are not labelled by *srp-GMA.* N.b. *VT62766-GAL4* image contrast enhanced to different parameters compared to other images owing to the very bright labelling of amnioserosal cells (cells on dorsal side of embryo destined to be removed during dorsal closure; labelled with an asterisk) in the stage 14 image. (f) labelling of smaller numbers of plasmatocytes on the ventral midline at stage 15 using *VT-GAL4* lines indicated and *UAS-GFP* (green); boxed regions show zooms of *VT-GAL4* positive plasmatocytes (f’). (g) ventral view of positive control embryo (*w;srp-GAL4/+;UAS-GFP*) and example plasmatocyte (g’) at stage 15. (h) scatterplot showing quantification of numbers of *VT-GAL4,UAS-GFP* labelled plasmatocytes on the ventral midline at stage 15; lines and error bars represent mean and standard deviation, respectively. P-values calculated via one-way ANOVA with a Dunnett’s multiple comparison post-test (all compared to *srp-GAL4* control; n=9 embryos per genotype. All scale bars represent 10μm. See Supplementary Table 1 for full list of genotypes.

### Subpopulations of *Drosophila* plasmatocytes vary across development: subpopulation dynamics in larvae and white pre-pupae

Having identified subpopulations of plasmatocytes in the embryo, we then tested other stages of the life cycle to see how expression might be maintained or modulated throughout development. In order to exclude potential expression in non-hemocyte cells (e.g. non-plasmatocyte cells apparent in Figure 2e), we labelled subpopulation cells specifically using a split GAL4 approach [31], via which only cells expressing both *serpent* (a well-characterised hemocyte marker; [32]) and the named VT enhancer would be labelled via transcriptional activation of *UAS* transgenes (Supplementary Figure 1).

Imaging of L3 larvae containing split GAL4 constructs (*srp-AD;VT-DBD* – henceforth abbreviated to *VTn*) driving *UAS-stinger* revealed that very few subpopulation cells were present at this stage (Figure 3a-f). From larval development onwards, we cannot use cell morphology to discriminate between plasmatocytes and other hemocyte lineages (crystal cells and lamellocytes) and therefore refer to subpopulation cells as hemocytes for these subsequent stages. Using this approach, *VT32897* and *VT17559* labelled the most cells (Figure 3c-d), with only the occasional cell present in *VT57089* larvae (Figure 3e) and cells essentially absent from *VT62766* larvae (Figure 3f). Labelled cells were also present in the head region, along the dorsal vessel (the fly heart) and between the salivary glands (which exhibit non-specific labelling) in *VT32897* larvae. The *VT32897* head region cells are potentially specifically localised hemocytes, whereas cells at the remaining two sites are likely to correspond to *serpent*-positive nephrocytes and garland cells [33,34], respectively (Figure 3d). *VT57089* shows additional staining in the head region (potentially the Bolwig organ; Figure 3e) and, as per the dorsal vessel-associated cells in *VT32897* (Figure 3d), this can be observed in positive controls (data not shown). These patterns closely resemble patterns observed using the initial *VT-GAL4* reporters, albeit with more restricted labelling due to our split GAL4 approach (data not shown).

**Figure 3.**
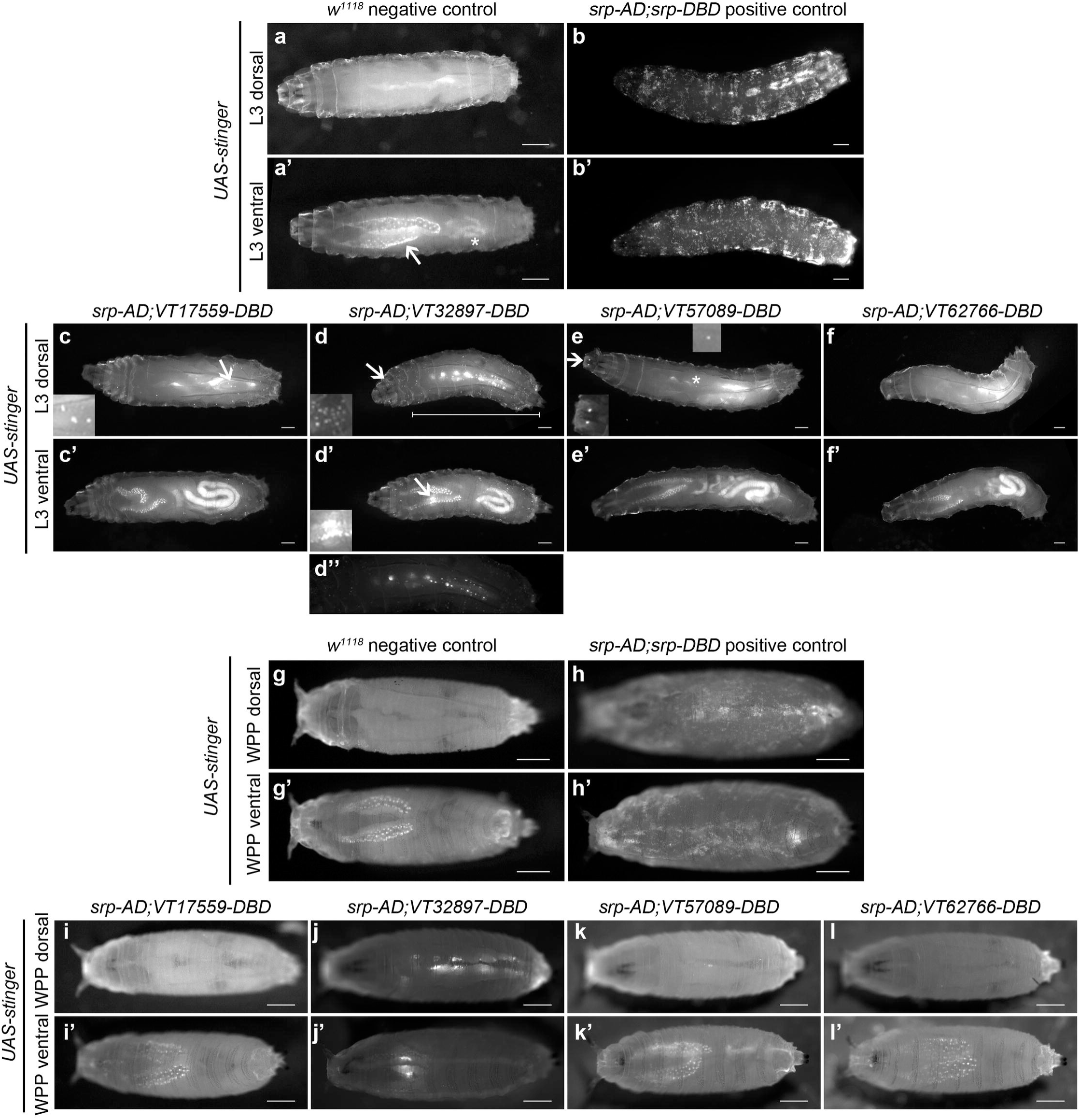
Plasmatocyte subpopulations in larvae and white pre-pupae. (a-f) dorsal and ventral views of L3 larvae lacking *GAL4* (a, negative control), with hemocytes labelled (b, positive control with *UAS-stinger* driven by *srp-AD;srp-DBD*), or with cells labelled through expression of *UAS-stinger* via *srp-AD* and the *VT-DBD* transgenes indicated (c-f); non-specific expression of Stinger in salivary glands and gut autofluorescence is visible in (a’) and (c’-f’) and is indicated by an arrow and asterisk, respectively, in (a’). Arrows/asterisks indicate regions containing circulating hemocytes (c), potential hemocyte population in the head region (d), possible proventricular region/garland cells (d’), cells in the Bolwig organ (arrow in e), and rare circulating hemocytes labelled via *srp-AD;VT57089-DBD* (asterisk in e) that are shown as zooms in inset images. (f”) shows region indicated by bar in (f) at a reduced brightness to reveal detail of cells along the dorsal vessel. (g-h) dorsal and ventral views of negative control (g, *UAS-stinger* but no driver) and positive control (h, *UAS-stinger* driven by *srp-AD;srp-DBD*) white pre-pupae (WPP). (i-l) VT enhancer-labelled cells are almost completely absent from WPP (*srp-AD;VT-DBD* transgenes used to drive *UAS-stinger* expression). All scale bars represent 500μm; images contrast enhanced to 0.3% saturation. See Supplementary Table 1 for full list of genotypes.

Imaging of white pre-pupae (WPP), the stage that marks the beginning of pupal development and metamorphosis, showed very similar patterns across the split GAL4 VT enhancer lines (Figure 3g-l), but with a further reduction in the numbers of cells labelled. It was possible to observe the occasional cell moving in circulation within WPP, strongly suggesting these cells are hemocytes (Supplementary Movies 2 and 3). Live imaging of *VT32897* WPP also confirmed association of cells with the pumping dorsal vessel (Supplementary Movie 4). Significantly, this data indicates that the presence of subpopulations within embryos is not simply a consequence of slow accumulation of fluorescent proteins by weak drivers, since these enhancer-based reporters do not label an ever-increasing number of cells as development proceeds. Overall, the numbers of hemocytes within subpopulations decreases over larval and early pupal stages demonstrating that plasmatocyte subpopulations are developmentally regulated. Such changes could reflect specific and changing requirements for specialised plasmatocyte subpopulations across the life cycle, for example an association with processes required for organogenesis [35–37]. This specific localisation of subpopulation cells also indicates the potential for tissue-resident macrophages in *Drosophila*.

### Subpopulation cells return in large numbers during pupal development

Since subpopulation cells appear associated with stages of development when organogenesis and tissue remodelling occurs, we hypothesised that hemocyte subpopulations would return during metamorphosis. Imaging pupae at various times after puparium formation (APF) revealed that subpopulation cells re-emerged in large numbers during this stage, but with distinct dynamics (Figure 4a-f): *VT17559* cells have already returned in very substantial numbers by 18h APF (Figure 4c), whereas *VT32897* reporter expression reappeared between 24 and 48h APF (Figure 4d). *VT57089* and *VT62766*-labelled cells increased in numbers more gradually over the course of pupal development (Figure 4e-f).

**Figure 4.**
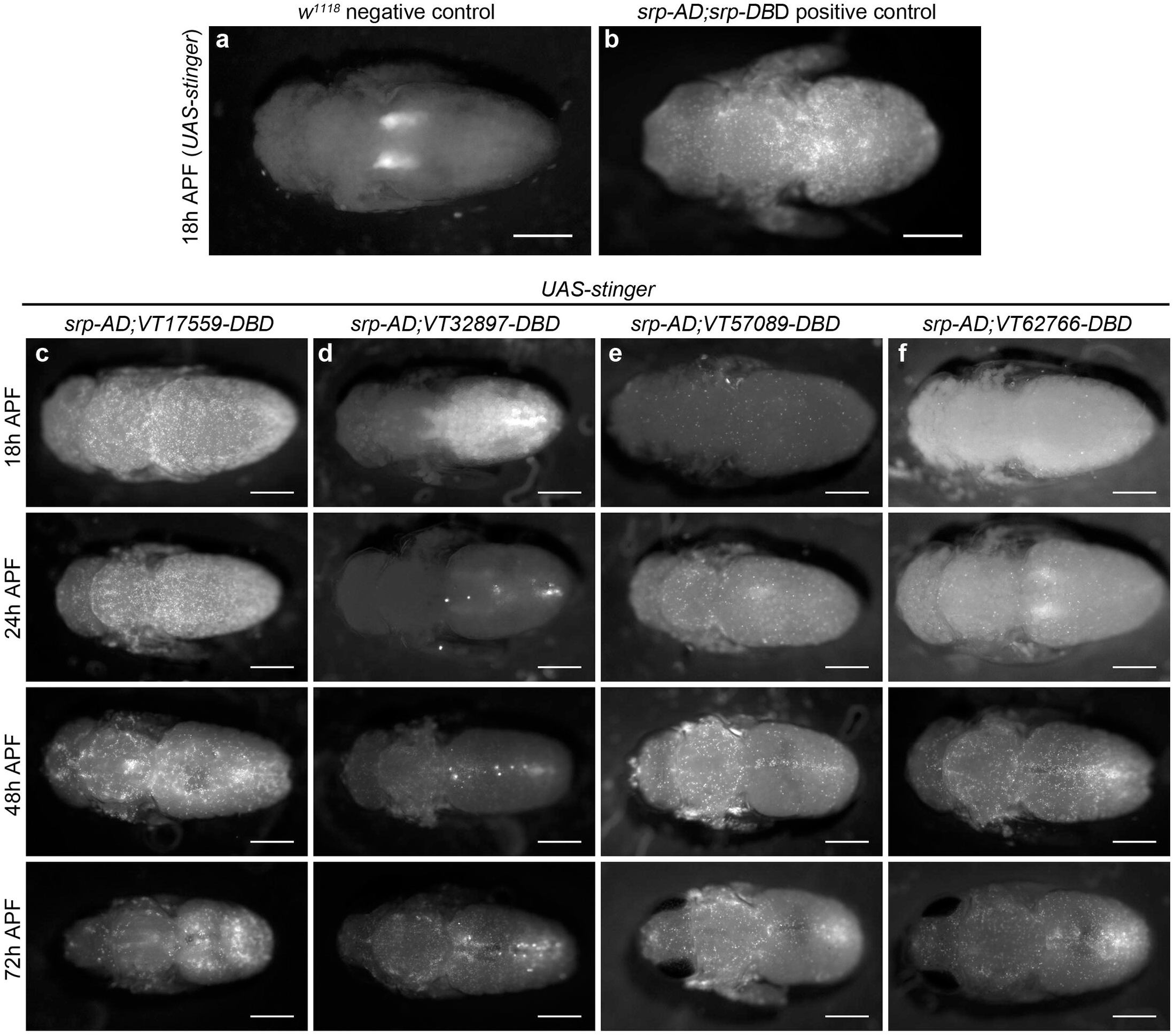
Plasmatocyte subpopulations return with distinct dynamics during pupal development. (a-b) dorsal images of negative control (lacking *GAL4* drivers, a) and positive control pupae (labelled via *srp-AD;srp-DBD*, b) at 18h after puparium formation (APF). (c-f) dorsal images showing localisation of cells labelled using *srp-AD* and *VT-DBD* (VT enhancers used to drive *DBD* expression indicated above panels) to drive expression of *UAS-stinger* during pupal development from 18h AFP to 72h APF (c-f). All image panels contrast enhanced to 0.3% saturation to reveal localisation of labelled cells due to differing intensities of reporter line expression. Scale bars represent 500μm. See Supplementary Table 1 for full list of genotypes.

### Subpopulations display distinct dynamics and localisation in adults

Immediately after adults hatch, large numbers of split GAL4-labelled cells can be observed across all lines and are present in selective regions that overlap with the overall adult hemocyte population (Figure 5a-e). The overall hemocyte population remains detectible as adults age (0-6 weeks; Figure 5a), however not all subpopulations exhibit an identical localisation or dynamics during this time (Figure 5b-e). *VT57089* and *VT62766* cells largely disappear by 1 week (Figure 5d-e) and the majority of *VT17559*-labelled cells are absent by 2 weeks (Figure 5b). By contrast, *VT32897* cells persist for at least 6 weeks of adult life and are particularly prominent in the thorax at 4 weeks (Figure 5c). Other differences in localisation are also apparent with cells particularly obvious in the legs for the *VT17559* line (Figure 5b, day 1-2 weeks), whereas *VT57089* and *VT62766*-labelled cells are more closely associated with the thorax and dorsal abdomen (Figure 5d-e, day 1). Labelled cells are also present in the proboscis for several lines (Figure 5c-e). The distinct dynamics of subpopulation cells strongly suggests these subpopulations are at least partially distinct from each other and highlights their plasticity during development, with their presence, return and disappearance correlating with changes in the biology of blood cells over the entire lifecourse.

**Figure 5.**
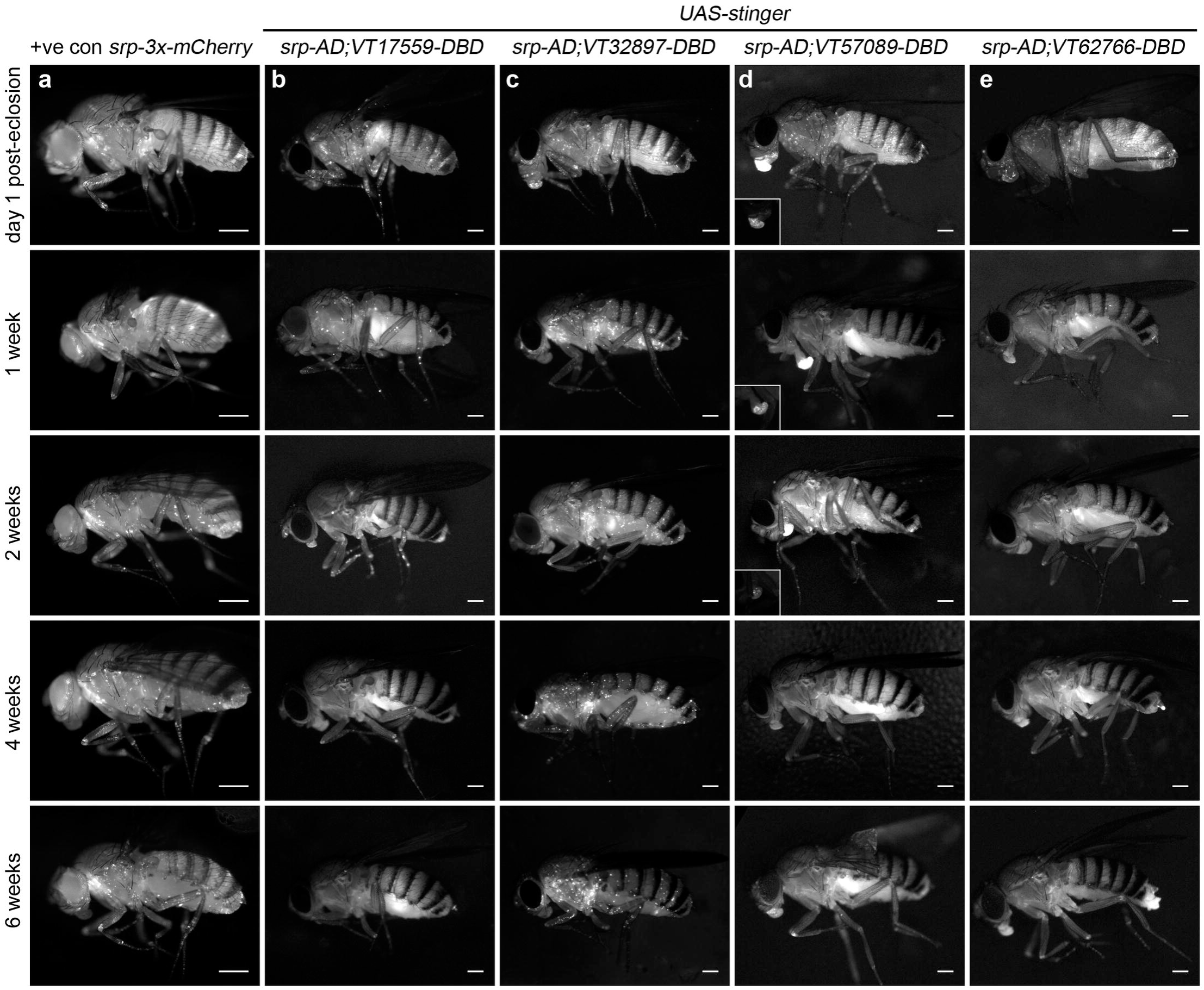
Plasmatocyte subpopulations exhibit distinct localisation and dynamics as adults age. (a-e) representative lateral images of adult flies between 0 and 6 weeks of age showing localisation of cells labelled using *srp-3x-mCherry* (positive control, a), or split GAL4 to drive expression of stinger (*srp-AD;VT-DBD*, b-e). The VT enhancers used to drive expression of the DNA binding domain (*DBD*) of *GAL4* correspond to *VT17559* (b), *VT32897* (c), *VT57089* (d) and *VT62766* (e); inset images show proboscis region at a reduced level of brightness to reveal cellular detail (d). Images contrast enhanced to 0.15% saturation (a-c, e) or 0.75% (d) to reveal localisation of labelled cells due to differing intensities of reporter line expression. Scale bars represent 500μm. At least 5 flies were imaged for each timepoint. See Supplementary Table 1 for full list of genotypes.

### Subpopulation cells behave in a functionally-distinct manner compared to the overall plasmatocyte population

Given that the VT lines identified above are specifically and dynamically expressed in subpopulations of hemocytes during *Drosophila* development, we next set out to investigate whether the labelled subpopulations are also functionally distinct using a range of immune-relevant assays. The ability of vertebrate macrophages to respond to pro-inflammatory stimuli, such as injuries, can vary according to their activation status [38,39]. To investigate this in our system, a well-established assay of inflammatory migration [19] was employed (Figure 1a; Supplementary Movie 1). Strikingly, following laser-induced wounding, cells labelled by three *VT-GAL4* lines (*VT17559-GAL4, VT32897-GAL4* and *VT62766-GAL4*) showed a significantly more potent migratory response to injury. In each case a greater proportion of labelled subpopulation cells migrated to wounds, compared to the overall hemocyte population as labelled by a pan-plasmatocyte driver (Figure 6a-c). Consistent with our results above, plasmatocytes labelled by the VT lines represent a subset of the total number of hemocytes present ventrally in stage 15 embryos (Figure 6d).

**Figure 6.**
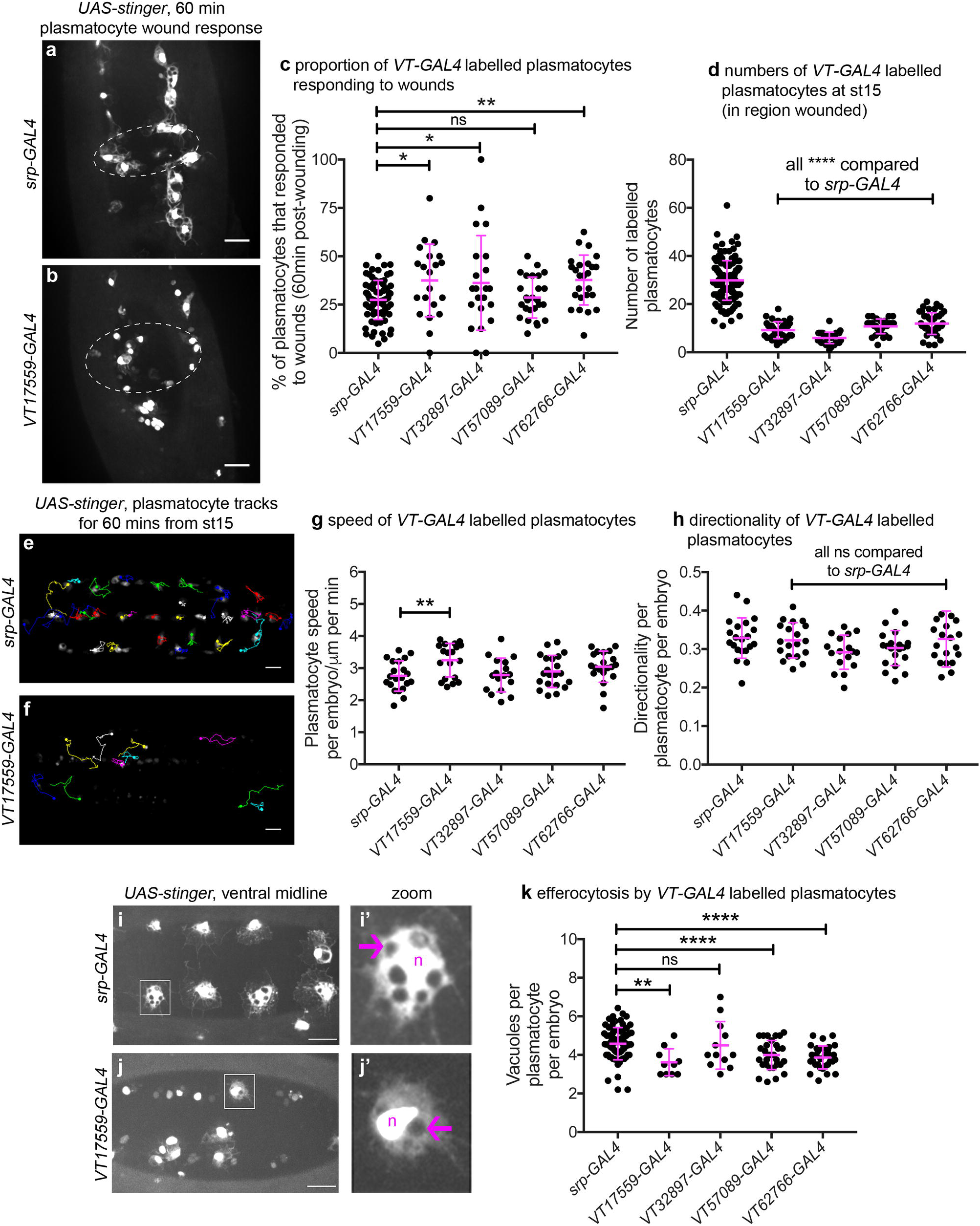
*Drosophila* plasmatocyte subpopulations demonstrate functional differences compared to the overall plasmatocyte population. (a-b) example images showing plasmatocyte wound responses at 60-minutes post-wounding (maximum projection of 15μm deep region). Cells labelled via *UAS-stinger* using *srp-GAL4* (a) and *VT17559-GAL4* (b); dotted lines show wound edges. (c-d) scatterplots showing percentage of *srp-GAL4* (control) or *VT-GAL4* labelled plasmatocytes responding to wounds at 60 minutes (c) or total numbers of labelled plasmatocytes in wounded region (d); p=0.018, 0.041, 0.99, 0.0075 compared to *srp-GAL4* (n=77, 21, 22, 26, 25) (c); p<0.0001 compared to *srp-GAL4* for all lines (n=139, 35, 37, 30, 44) (d). (e-f) example tracks of plasmatocytes labelled with GFP via *srp-GAL4* (e) and *VT17559-GAL4* (f) during random migration on the ventral side of the embryo for 1 hour at stage 15. (g-h) scatterplots showing speed per plasmatocyte, per embryo (g) and directionality (h) at stage 15 in embryos containing cells labelled via *srp-GAL4* (control) or the *VT-GAL4* lines indicated; p=0.0097, 0.999, 0.82, 0.226 compared to *srp-GAL4* (n=21, 19, 17, 21, 20) (g); p=0.998, 0.216, 0.480, 0.999 compared to *srp-GAL4* (n=21, 19, 17, 21, 20) (h). (i-j) example images of cells on the ventral midline at stage 15 with labelling via *UAS-stinger* expression using *srp-GAL4* (i) and *VT17559-GAL4* (j); zoomed plasmatocytes (i’, j’) indicated by white boxes in main panels; arrows show vacuoles, “n” marks nucleus; n.b. panels contrast enhanced independently to show plasmatocyte morphology. (k) scatterplots showing vacuoles per plasmatocyte, per embryo at stage 15 (measure of efferocytosis/apoptotic cell clearance); cells labelled via *srp-GAL4* (control) or the *VT-GAL4* lines indicated; p=0.0020, 0.99, 0.0040, 0.0002 compared to *srp-GAL4* (n=76, 10, 12, 29, 31). Lines and error bars represent mean and standard deviation, respectively (all scatterplots); one-way ANOVA with a Dunnett’s multiple comparison test used to compare *VT-GAL4* lines with *srp-GAL4* control in all datasets; ns, *, ** and **** denote p>0.05, p<0.05, p<0.01 and p<0.0001, respectively. All scale bars represent 20μm. See Supplementary Table 1 for full list of genotypes.

We next investigated in vivo migration speeds of the embryonic plasmatocyte subpopulations (as per Figure 1c-d). Stage 15 embryos were imaged for 1 hour and individual plasmatocyte movements were tracked (Figure 6e-f). Only the *VT17559-GAL4* labelled plasmatocyte subpopulation displayed statistically significantly faster rates of migration compared to the overall plasmatocyte population (labelled using *srp-Gal4*; Figure 6g). There were no differences in directionality (cell displacement divided by total path length) for any of the subpopulations, suggesting that the mode of migration was similar across these lines and with that of the overall population (Figure 6h).

Apoptotic cell clearance (efferocytosis) represents another evolutionarily-conserved function performed by embryonic plasmatocytes (Figure 1e; [40]). Therefore, we investigated this function in subpopulations, using numbers of vacuoles per cell as a proxy for this process [18]. Cells labelled via *VT17559-GAL4, VT57089-GAL4* and *VT62766-GAL4* (but not *VT32897-GAL4*) contained fewer vacuoles than the overall plasmatocyte population (Figure 6i-k), suggesting that these discrete populations of cells are less effective at removing apoptotic cells inside the developing embryo.

Finally, we examined cell size and shape of labelled plasmatocyte subpopulations. Vertebrate macrophages are highly heterogeneous, with distinct morphologies dependent upon their tissue of residence or polarisation status [41–43]. We found no obvious size or shape differences between *VT-GAL4* labelled cells and the overall plasmatocyte population (Supplementary Figure 2a-e). This was also the case when *VT-GAL4* positive cells were compared to internal controls (*VT-GAL4* negative cells within the same embryos) for a range of shape descriptors (Supplementary Figure 2f-i). Similarly, we were unable to detect differences in ROS levels or the proportion of *VT-GAL4* labelled plasmatocytes that phagocytosed pHrodo-labelled *E. coli* compared to controls (Supplementary Figures 3 and 4), two processes associated with pro-inflammatory activation of macrophages [44].

Taken together these data show that the subpopulations of plasmatocytes identified via the *VT-GAL4* reporters exhibit functional differences compared to the overall plasmatocyte population (Table 1). Therefore, as well as displaying molecular differences in the form of differential enhancer activity, and hence reporter expression, these discrete populations of cells behave differently. This strongly suggests that these cells represent functionally-distinct subpopulations and that the plasmatocyte lineage is not homogeneous. Furthermore, not all subpopulations displayed identical functional characteristics, suggesting that there are multiple distinct subtypes present in vivo, although some overlap between subpopulations seems likely. For example, *VT17559-GAL4* labelled cells were more effective at responding to wounds and migrated more rapidly, but carried out less phagocytosis of apoptotic cells. By contrast, *VT32987-GAL4* labelled cells only displayed improved wound responses (Figure 6).

**Table 1.** Summary of plasmatocyte subpopulation characteristics and their developmental regulation.

|  |  |  |  |  |  |  |  |  |  |  |
| --- | --- | --- | --- | --- | --- | --- | --- | --- | --- | --- |
|  | Subpopulation characteristics (compared to overall population): |  |  |  |  | Subpopulations in: |  |  |  |  |
| Subpopulation | Wound responses | Migration speed | Efferocytosis | ROS levels | Phagocytosis of *E. coli* | Embryos | Larvae | Pupae | Newly hatched adults | Aged adults |
| **VT17559** | ↑ | ↑ | ↓ | no difference | no difference | distinct subpopulation | very few cells labelled | large numbers labelled by 18h APF | large numbers present | largely absent by 2 weeks |
| **VT32897** | ↑ | no difference | no difference | no difference | no difference | distinct subpopulation (fewest cells) | few cells labelled + nephrocytes & garland cells (?) | large numbers labelled by 72h APF | large numbers present | labelled cells persist |
| **VT57089** | no difference | no difference | ↓ | no difference | no difference | distinct subpopulation | almost no cells labelled + Bolwig Organ (?) | steady increase in numbers labelled | large numbers present | largely absent by 1 week |
| **VT62766** | ↑ | no difference | ↓ | no difference | no difference | distinct subpopulation | almost no cells labelled | large numbers labelled by 48h APF | large numbers present | largely absent by 1 week |

### VT enhancers identify functionally active genes within plasmatocytes

In the original study that generated the *VT-GAL4* collection, the majority of active enhancer fragments tested were found to control transcription of neighbouring genes [29]. Thus, genes proximal to enhancers that label plasmatocyte subpopulations represent candidate regulators of immune cell function (Table 2; Figure 7a). *VT62766-GAL4* labels a subpopulation of plasmatocytes with enhanced migratory responses to injury (Figure 6) and this enhancer region is found within the genomic interval containing *paralytic* (*para*), which encodes a subunit of a voltage-gated sodium channel [45], and upstream of the 3’ end of *calnexin14D* (*cnx14D*) (Figure 7a). *cnx14D* encodes a calcium-binding chaperone protein resident in the endoplasmic reticulum [46]. Alterations in calcium dynamics are associated with clearance of apoptotic cells [47,48] and modulating calcium signalling within plasmatocytes alters their ability to respond to wounds [49]. Therefore, given the association of *cnx14D* with the *VT62766* enhancer and the potential for plasmatocyte behaviours to be modulated by altered calcium dynamics, we examined whether misexpressing *cnx14D* in all plasmatocytes was sufficient to cause these cells to behave more similarly to the *VT62766* subpopulation. Critically, pan-hemocyte expression of *cnx14D* stimulated wound responses with elevated numbers of plasmatocytes responding to injury compared to controls (Figure 7b-c), consistent with the enhanced wound responses of the endogenous *VT62766-GAL4* positive plasmatocyte subpopulation (Figure 6c). This reveals that genes proximal to subpopulation-defining enhancers represent candidate genes in dictating the biology of cells in those subpopulations. More importantly, misexpression of a subpopulation-linked gene promotes a similar behaviour to that subpopulation in the wider plasmatocyte population.

**Table 2.** VT enhancer region location and neighbouring genes.

| **VT enhancer** | **Genomic region\*** | **Nearest genes** | **Distance of enhancer from gene** |
| --- | --- | --- | --- |
| *VT17559* | chr2R 12069698–12070780 | *Lis-1*<br>*CG8441*<br>*Ptp52F* | overlapping<br>2929bp upstream<br>3887bp downstream |
| *VT32897* | chr3L 18631149–18633281 | *MYPT-75D*<br>*bora*<br>*not* | overlapping<br>13299bp downstream<br>15921bp downstream |
| *VT57089* | chrX 4961770–4962316 | *ovo*<br>*CG32767*<br>*CR44833* | overlapping<br>3290bp upstream<br>3870bp downstream |
| *VT62766* | chrX 16406666–16408777 | *para*<br>*cnx14D*<br>*CG9903* | overlapping<br>10404bp upstream<br>26520bp upstream |

\* *D. melanogaster* Apr. 2006 (BDGP R5/dm3) Assembly
Data taken from http://enhancers.starklab.org/

**Figure 7.**
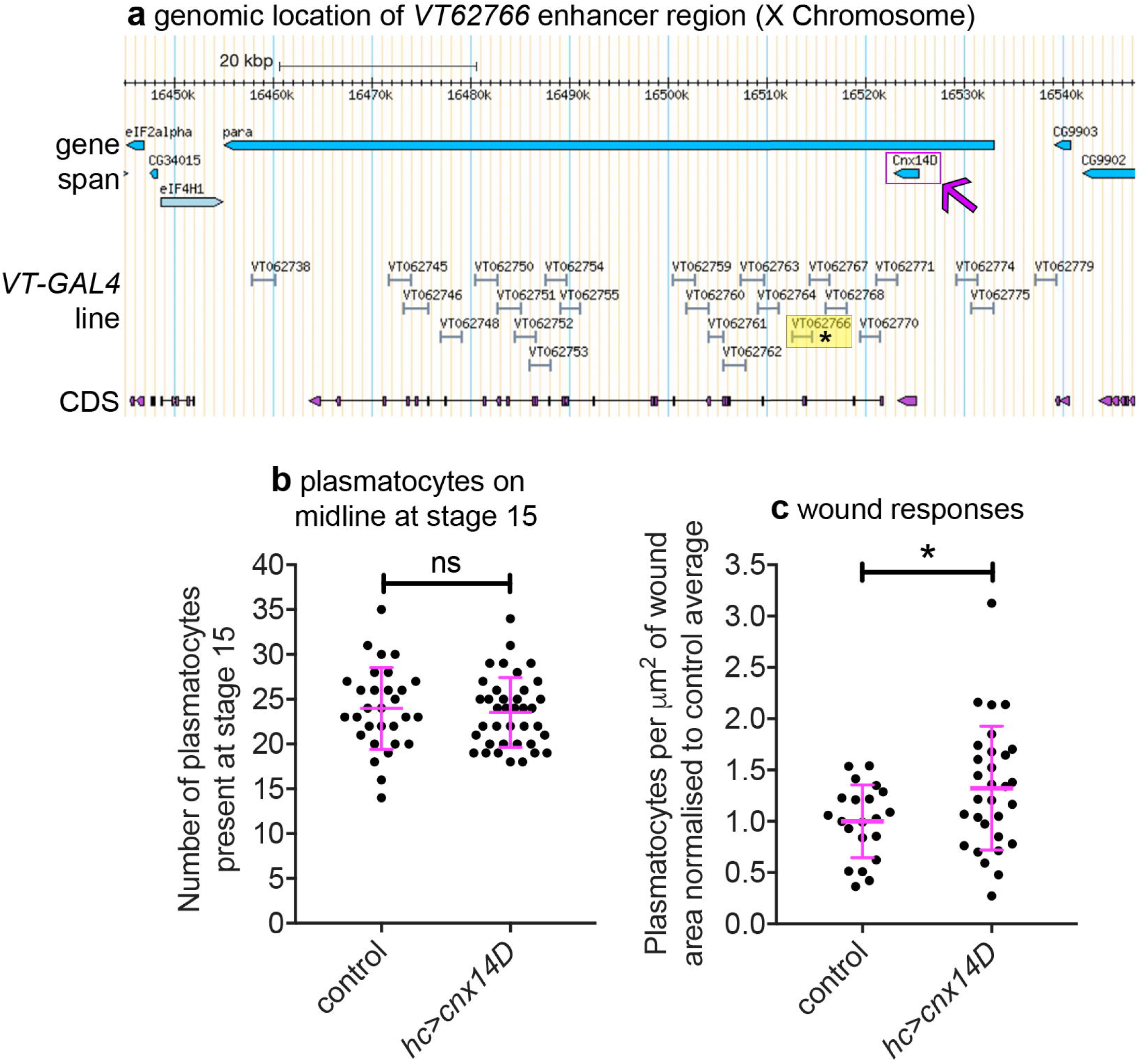
Misexpression of *cnx14D* improves plasmatocyte inflammatory responses to injury. (a) chromosomal location of the *VT62766-GAL4* enhancer region. Only one transcript is shown for *para*, which possesses multiple splice variants. The *VT62766* region is highlighted in yellow and by an asterisk; *cnx14D* (indicated by magenta arrow) lies within *para*. (b) scatterplot showing numbers of plasmatocytes present at stage 15 on the ventral side of the embryo ahead of wounding in controls and on misexpression of *cnx14D* in all hemocytes using both *srp-GAL4* and *crq-GAL4* (*hc>cnx14D*); n=30 and 38 for control and *hc>cnx14D* embryos, respectively, p=0.670 via Student’s t-test. (c) scatterplot of wound responses 60-minutes post-wounding (number of plasmatocytes at wound, normalised for wound area and to control responses); n= 21 and 30 for control and *hc>cnx14D* embryos, respectively; p=0.0328 via Student’s t-test. Line and error bars represent mean and standard deviation, respectively (b-c). See Supplementary Table 1 for full list of genotypes.

### Plasmatocyte subpopulations can be modulated via exposure to enhanced levels of apoptosis

Having defined functional differences in embryonic plasmatocyte subpopulations and characterised how these populations shift during development and ageing, we sought to identify the processes via which these subpopulations were specified. In vertebrates, a range of stimuli drive macrophage heterogeneity and polarisation [3,4], with apoptotic cells able to polarise macrophages towards anti-inflammatory phenotypes [50,51]. In the developing fly embryo, high apoptotic cell burdens impair wound responses [52,53], consistent with reprogramming of plasmatocytes towards less wound-responsive states. In order to test whether apoptotic cells might regulate plasmatocyte subpopulations, we exposed plasmatocytes to increased levels of apoptosis in vivo. In the developing fly embryo, both glial cells and plasmatocytes contribute to the clearance of apoptotic cells. We, and others, have previously shown that loss of *repo*, a transcription factor required for glial specification [54–56], leads to decreased apoptotic cell clearance by glia [57], and a subsequent challenge of plasmatocytes with increased levels of developmental apoptosis (Figure 8a-b; [53]). Therefore, a *repo* mutant background represents an established model with which to stimulate plasmatocytes with enhanced levels of apoptosis.

**Figure 8.**
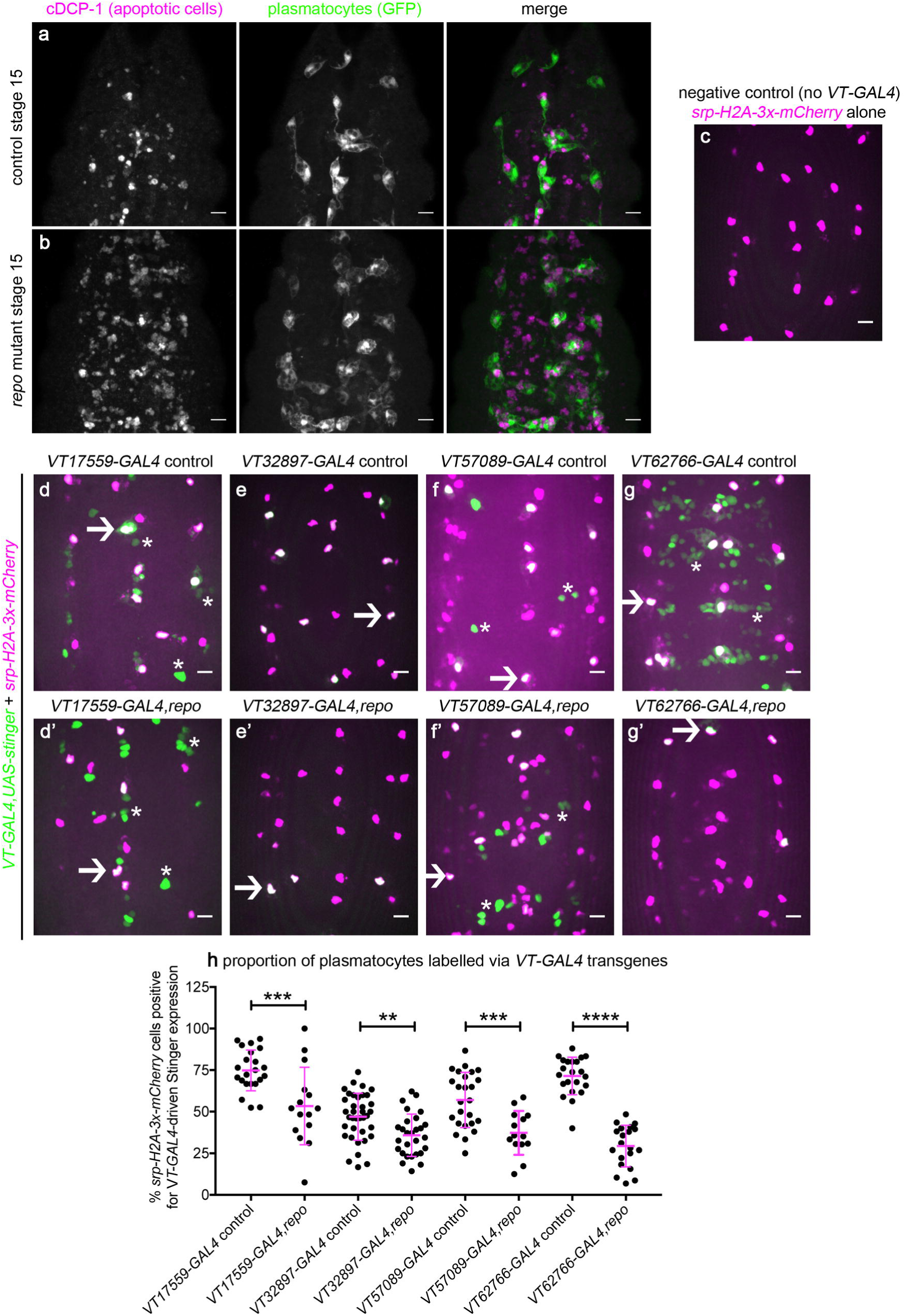
*Drosophila* plasmatocyte subpopulation identity can be controlled through exposure to apoptotic cells. (a-b) maximum projections showing apoptotic cells (via anti-cDCP-1 staining, magenta in merge) and plasmatocytes (via anti-GFP staining, green in merge) at stage 15 on the ventral midline in control and *repo* mutant embryos. (c-g) maximum projections of the ventral midline showing a negative control embryo (c) and embryos containing *VT-GAL4* labelled plasmatocytes at stage 15 in control (d-g) and *repo* mutant embryos (d’-g’). *VT-GAL4* used to drive *UAS-stinger* expression (green) and *srp-H2A-3x-mCherry* used to label plasmatocytes (magenta). Arrows and asterisks indicate examples of *VT-GAL4* positive plasmatocytes and non-plasmatocyte cells, respectively; note loss of non-plasmatocyte *VT-GAL4* expression in *repo* mutants versus controls for *VT62766-GAL4*. (h) scatterplot showing percentage of H2A-3x-mCherry positive cells that are also positive for *VT-GAL4* driven Stinger expression in control and *repo* mutant embryos at stage 15. Student’s t-test used to show significant difference between controls and *repo* mutants (p=0.0009, n=22, 15 for *VT17559-GAL4* lines; p=0.0017, n=37, 28 for *VT32897-GAL4* lines; p=0.0005, n=25, 14 for *VT57089-GAL4* lines; p<0.0001, n=22, 20 for *VT62766-GAL4* lines). Scale bars represent 10μm (a-g); lines and error bars represent mean and standard deviation (h); **, *** and **** denote p<0.01, p<0.001 and p<0.0001, respectively. See Supplementary Table 1 for full list of genotypes.

Using *srp-H2A-mCherry* to mark all plasmatocytes within the embryo (Figure 8c), we quantified the proportion of plasmatocytes labelled via *VT-GAL4* transgenes in *repo* mutants compared to controls (Figure 8d-h). Increased exposure to apoptotic death shifted plasmatocytes out of each subpopulation (Figure 8d-h). Subpopulations exhibited differing sensitivities to contact to apoptotic cells, with *VT62766-GAL4* labelled cells undergoing the largest decrease in labelled cells in a *repo* mutant background (Figure 8h). These results therefore reveal a mechanism via which the molecularly and functionally-distinct subpopulations of plasmatocytes we have identified can be manipulated using an evolutionarily-conserved, physiological stimulus (apoptotic cells) relevant to immune cell programming.

## Discussion

We have identified molecularly and functionally-distinct subpopulations of *Drosophila* macrophages (plasmatocytes). These subpopulations showed functional differences compared to the overall plasmatocyte population, exhibiting enhanced responses to injury, faster migration rates and reduced rates of apoptotic cell clearance within the developing embryo. These subpopulations are highly plastic with their numbers varying across development, in line with the changing behaviours of *Drosophila* blood cells across the lifecourse. That these discrete populations of plasmatocytes represent bona fide subpopulations is evidenced by the finding that numbers of cells within subpopulations can be manipulated via exposure to enhanced levels of apoptotic cell death in vivo. Furthermore, pan-hemocyte expression of a gene (*cnx14D*) linked to one of the enhancers used to visualise these subpopulations (*VT62766-GAL4*) shifts the behaviour of these cells towards a more wound-responsive state, resembling the behaviour of *VT62766-GAL4* labelled cells. Taken together this data strongly suggests that *Drosophila* blood cell lineages are more complex than previously known.

Vertebrate macrophage lineages show considerable heterogeneity due to the presence of circulating monocytes, a wide variety of tissue resident macrophages and a spectrum of activation states that can be achieved. Whether more simple organisms such as *Drosophila* exhibit heterogeneity within their macrophage-like lineages has been a topic of much discussion and hints in the literature suggest this as a possibility. The ease of extracting larval hemocytes has meant these cells have received more attention than their embryonic counterparts. Braun and colleagues identified heterogeneity in reporter expression within plasmatocytes in an enhancer trap screen, but without associating these with functional differences [58]. Non-uniform expression has also been reported for plasmatocyte genes such as *hemolectin* [59], *hemese, nimrod* [60,61], *croquemort* [26], TGF-β family members [26] and the iron transporter *malvolio* [62], though some of these differences are likely due to incomplete differentiation from a pro-hemocyte state [25]. Recent transcriptional profiling approaches via scRNA-seq have suggested the existence of distinct larval blood cell populations in *Drosophila* [27,28]. One study interpreted this data as reflecting different progenitor/differentiation states [27]; another identified a number of potentially different functional groups, including more activated cell populations displaying expression signatures reflective of active Toll and JNK signalling [28]. Our identification of developmentally-regulated subpopulations, coupled with this recent evidence from larvae, points to heterogeneity within the plasmatocyte lineage.

The subpopulations we have identified are almost entirely absent from L3 larvae (the stage used in the aforementioned scRNA-seq studies) and presumably represent additional heterogeneity specific to other developmental stages. It is clear that the biology of *Drosophila* blood cells varies significantly across the lifecourse: for instance plasmatocytes play strikingly different functional roles in embryos and larvae [35,36], shifting from developmental roles to host defence. Additionally, modes of migration to sites of injury are similar in embryos and pupae (directional migration [19,63]), but larval cells are captured from circulation via adhesion [64]. These functional differences are reflected in molecular differences between embryonic and larval blood cells revealed via bulk RNA-seq [28], with reprogramming within larvae potentially explaining why our VT enhancer-labelled subpopulations are absent at that stage. Transcriptional changes are also associated with steroid hormone-mediated signalling in pupae [37], which may drive re-emergence of subpopulations in time for metamorphosis.

In higher vertebrates, erythro-myeloid precursor/progenitor cells seed the developing embryo to give rise to tissue resident macrophage populations [65–67]. Intriguingly, the localisation of subpopulations in adult flies shows some biases between subpopulation lines and the overall population, hinting at the potential for some degree of tissue residency in *Drosophila*. Hemocytes localise to and/or play specialised roles at a range of tissues including the respiratory epithelia [68], dorsal vessel [69], ovaries [70], wings [71], gut [72] and proventriculus [73]. It is therefore tempting to speculate that particular subpopulations could be recruited or differentiate in situ in order to carry out specific functions.

Macrophage diversity enables these important innate immune cells to operate in a variety of niches and carry out a wide variety of functions in vertebrates. Our data demonstrate that not all macrophages are equivalent within the developing *Drosophila* embryo, although the enhancers we have used to identify plasmatocyte subpopulations do not correspond to markers used in defining macrophage polarisation or tissue resident populations in an obvious way. Therefore how the subpopulations we have uncovered map onto existing vertebrate paradigms remains an open question. Nonetheless, the subpopulations we have identified could be viewed as a displaying a pro-inflammatory skewing of immune cell behaviours, given their enhanced wound responses, faster rates of migration and decreased efferocytic capacity. Pro-inflammatory macrophages (M1-like) in vertebrates are associated with clearance of pathogens, release of pro-inflammatory cytokines and, most pertinently, initial responses to injury [44]. In contrast, anti-inflammatory macrophages (M2-like) are more allied with tissue development and repair [74] and can display enhanced rates of efferocytosis [75–77].

Apoptotic cell clearance can promote anti-inflammatory states in vertebrates [78]. Consequently, it is both consistent and compelling that exposure of *Drosophila* plasmatocytes to excessive levels of apoptotic cells dampens their inflammatory responses to injury and rates of migration in the developing embryo [18,52,53] and also shifts cells out of the more wound-responsive and potentially pro-inflammatory subpopulations we have discovered. Other precedents may be apparent in flies with shifts towards aerobic glycolysis occurring during infection [79], similar to those observed in vertebrate polarisation to pro-inflammatory states [80]. Furthermore, TGF-β signalling is associated with promotion of anti-inflammatory characteristics in vertebrates during resolution of inflammation [78] and these molecules can be found in discrete sets of hemocytes on injury and infection in adult flies [26]. Thus, despite significant evolutionary distance between flies and vertebrates, comparable processes and mechanisms may control the behaviours of their innate immune cells.

We have concentrated on using the VT enhancers as reporters to follow subpopulation behaviour in vivo, however these elements also potentially identify genes required for specific functions associated with each subpopulation. For instance, the *VT17559* enhancer overlaps *Lisencephaly-1*, which has been shown to be expressed in hemocytes [81]. Furthermore, misexpression of *cnx14D*, located proximally to the *VT62766* enhancer, was sufficient to improve overall wound responses, paralleling the behaviour of the *VT62766-GAL4* labelled subpopulation. Cnx14D can bind calcium and therefore potentially modulates calcium signalling within plasmatocytes. Calcium signalling is known to influence wound responses in flies [49] and plays a central role during phagocytosis of apoptotic cells [47,48]. Therefore a molecule such as Cnx14D, which also has a known role in phagocytosis in *Dictyostelium* [82], could help fine-tune the behaviour of specific macrophage subpopulations. When considered in combination with the ability to manipulate the numbers of cells within subpopulations with physiologically relevant stimuli, the functional linkage of candidate genes with subpopulation behaviours strongly suggests that we have identified bona fide functionally and molecularly-distinct macrophage subpopulations in the fly.

In conclusion, we have demonstrated that *Drosophila* macrophages are a heterogeneous population of cells with distinct functional capabilities. We have characterised novel tools in which to visualise these subpopulations and have used these tools to reveal functional differences between these subpopulations and the general complement of hemocytes. Furthermore, we have shown that these subpopulations can be manipulated by exposure to apoptotic cells and can be linked to specific functional players. Therefore, we have further established *Drosophila* as a model for studying macrophage heterogeneity and immune programming and demonstrate that macrophage heterogeneity is a key feature of the innate immune system even in the absence of adaptive immunity and is conserved more widely across evolution than previously anticipated.

## Methods

### Fly genetics and reagents

Standard cornmeal/agar/molasses media was used to culture *Drosophila* at 25°C (see Supplementary Table 2 for ingredients). *srp-GAL4* [83], *crq-GAL4* [19], *da-GAL4* [84] and the *GAL4*-independent lines *srp-GMA* [30], *srp-3x-mCherry* and *srp-H2A-3xmCherry* [85] were used to label the entire hemocyte population during embryonic development or in adults. *Hml(Δ)-GAL4* was used to label larval hemocytes [86]. *srp-GAL4, Hml(Δ)-GAL4, VT-GAL4* lines (obtained from the VDRC, Vienna; [29]) and split GAL4 lines (see below) were used to drive expression from *UAS-tdTomato* (Bloomington stock 36327), *UAS-GFP, UAS-red stinger, UAS-stinger, UAS-cnx14D* (Harvard stock d04188) or *UAS-GC3ai* [87]. Experiments were conducted in a *w*^*1118*^ background and the *repo*^03702^ null allele was used to expose plasmatocytes to enhanced levels of apoptotic cell death in the embryo [53,54,56]. Both *UAS-tdTomato* and *UAS-GFP* were used to analyse subpopulations in the developing embryo in order to ensure labeling of discrete numbers of plasmatocytes was not due to positional effects of insertion sites that led to mosaic expression (Figure 2). See Supplementary Table 1 for a full list of *Drosophila* genotypes, transgenes and the sources of the *Drosophila* lines used in this study.

Flies were added to laying cages attached to apple juice agar plates supplemented with yeast paste and allowed to acclimatise for 2 days before embryo collection. Plates were then changed every evening and cages incubated at 22°C overnight before embryos were collected the following morning. Embryos were collected by washing the plates with distilled water and gently disturbing the embryos with a paintbrush, after which embryos were collected into a cell strainer. Embryos were dechorionated in undiluted bleach for 1-2 minutes and then washed in distilled water until free from bleach. The fluorescent balancers *CTG, CyO dfd, TTG* and *TM6b dfd* [88,89] were used to discriminate homozygous embryos after removal of the chorion.

### Generation of split GAL4 transgenic lines

We used the split GAL4 system [31] to restrict VT enhancer expression to *serpent*-positive cells. The activation domain (AD) of *GAL4* was expressed using a well-characterised fragment of the hemocyte-specific *serpent* promoter [83,85] and the DNA-binding domain (DBD) was expressed under the control of VT enhancer regions corresponding to *VT17559-GAL4, VT32897-GAL4, VT57089-GAL4* or *VT62766-GAL4*. High-fidelity polymerase (KAPA HiFi Hotstart ReadyMix, Roche) was used to PCR amplify VT enhancer regions from *w*^*1118*^ genomic DNA, which were then TA cloned into the pCR8/GW/TOPO vector. Primers were designed according to VT enhancer sequences available via the Stark Lab Fly Enhancers website (http://enhancers.starklab.org/; [29]). To make *VT-DBD* transgenic constructs, VT enhancers were transferred from *pCR8/GW/TOPO* into *pBPZpGal4DBDUw* (Addgene clone 26233) using LR clonase technology (Invitrogen Gateway LR Clonase II Enzyme Mix - Catalog Number 11791-020).

To express the *DBD* and *AD* of *GAL4* under the control of the *serpent* promoter (*srp-AD* and *srp-DBD*), these were subcloned into an attB containing vector containing this promoter (*pBS_MCS_SRPW_attB*; DSPL337 – a gift from Daria Siekhaus, IST, Austria; [85]). *DBD* and *AD* sequences along with the *Drosophila* synthetic minimal core promoter (DSCP) region were amplified using PCR from vectors *pBPZpGal4DBDUw* and *pBPp65ADZpUw* (Addgene clone 26234) using primers that added NotI and AvrII restriction sites (CTGATCGCGGCCGCAAAGTGGTGATAAACGGCCGGC and GATCAGCCTAGGGTGGATCTAAACGAGTTTTTAAGCAAACTCAC). These were subcloned into DSPL337 cut with NotI/AvrII (New England Biolabs) using T4 DNA ligase (Promega). Transgenic flies were generated by site-specific insertion of transgenic constructs into the VK1 attP site on chromosome 2 and/or attP2 on chromosome 3 by Genetivision (Texas, USA).

### Imaging of *Drosophila* embryos, larvae, pupae and adults

Live embryos were mounted ventral-side up on double-sided sticky tape in a minimal volume of Voltalef oil (VWR), after dechorionation in bleach as per Evans et al., 2010 [90]. High-resolution live imaging of plasmatocytes was carried out on an UltraView Spinning Disk system (Perkin Elmer) using a40x UplanSApo oil immersion objective lens (NA 1.3). A Nikon A1 confocal microscope was used to image plasmatocyte morphology (40x CFI Super Plan Fluor ELWD oil immersion objective lens, NA 0.6) and a Zeiss Airyscan microscope (40x Plan-Apochromat oil immersion objective lens, NA 1.4) was used for imaging of embryos stained with ROS dyes.

Wandering L3 Larvae were removed from straight-sided culture bottles containing the food on which they were reared at 25°C and cleaned in distilled water. Larvae were then imaged in fresh ice-cold, distilled water using a MZ205 FA fluorescent dissection microscope with a 2x PLANAPO objective lens (Leica) and LasX software (Leica). White pre-pupae were collected from the same culture bottles and washed before imaging on the same system, which was also used to image subsequent stages of development. For analysis of plasmatocyte populations in pupae, white pre-pupae were also collected, aged at 25°C and the pupal case removed at a range of times after puparium formation. Dissected pupae were covered with halocarbon oil 500 (Sigma-Aldrich) to prevent desiccation during imaging. For imaging of plasmatocyte populations in adults, females were aged in vials containing cornmeal/agar/molasses media at 25°C, with no more than 7 flies kept per vial. Flies were transferred to new food vials every 2-3 days.

### Wounding assay

Live stage 15 embryos were prepared and mounted as described above. The ventral epithelium of the embryos was ablated on the ventral midline using a Micropoint nitrogen-pulsed ablation laser (Andor) fitted to an Ultraview spinning disk confocal system (PerkinElmer) as as per Evans et al., 2015 [91]. Pre-wound z-stacks of 30μm were taken of superficial plasmatocytes with a 1μm z-spacing between z-slices. Post-wound images were taken on the same settings either at 2-minute intervals for 60 minutes (Figure 1) or at the end timepoint of 60 minutes (Figures 6 and 7).

The proportion of plasmatocytes labelled with *UAS-stinger* (expression via *srp-GAL4* or *VT-GAL4*) was assessed by counting the number of labelled cells at or in contact with the wound site within a 35μm deep volume on the ventral midline at 60-minutes post-wounding; this was divided by the total number of labelled cells present within the stack to calculate the percentage of plasmatocytes responding to injury. The brightfield channel was used to visualise the wound margin and only those embryos with wounds between 1000μm^2^ and 4000μm^2^ were included in analyses. Quantification was performed on blinded images in Fiji.

### Quantification of migration speeds/random migration

Embryos were prepared and mounted as described by Evans et al., 2010 [90]. Random migration was imaged using a spinning disk system (Ultraview, PerkinElmer), with an image taken every 2 minutes for 1 hour with a z-spacing of 1μm and approximately 20μm deep from the ventral nerve cord using a 20x UplanSApo air objective lens (NA 0.8). Maximum projections were made for each timepoint (25μm depth) and the centre of individual plasmatocyte cell bodies tracked using the manual tracking plugin in Fiji. Random migration speed (μm/min) and directionality (the ratio of the Cartesian distance to the actual distance migrated) were then calculated using the Ibidi chemotaxis plugin.

### Quantification of apoptotic cell clearance

The number of vacuoles per plasmatocyte (averaged per embryo) was used as a read-out of apoptotic cell clearance as per Evans et al., 2013 [18]. Vacuoles were counted using z-stacks of GFP-labelled plasmatocytes taken from live imaging experiments. Vacuoles were scored in the z-slice in which each macrophage exhibited its maximal cross-sectional area. Only labelled plasmatocytes present on the ventral midline of stage 15 embryos were included. Analysis was performed on blinded image stacks. This analysis does not report the absolute numbers of apoptotic corpses per cell but provides a relative read-out of the phagocytic index.

### Fixation and immunostaining of embryos

Embryos were fixed and stained as per Roddie et al., 2019 [52]. Embryos containing plasmatocytes labelled via *srp-GMA* and *GAL4*-driven tdTomato expression were fixed, then mounted in Dabco mountant. Control and *repo* mutant embryos containing plasmatocytes labelled via *crq-GAL4,UAS-GFP* were fixed and immunostained using mouse anti-GFP (ab1218 1:200; Abcam) and rabbit anti-cleaved DCP-1 (9578S 1:1000; Cell Signaling Technologies) to detect plasmatocytes and apoptotic cells, respectively. Embryos were imaged on the Nikon A1 system described above.

### Dissection, culture and stimulation of larval hemocytes

Hemocytes were dissected from wandering L3 larvae by ripping open larvae from the posterior end in S2 cell media, which consists of Schneider’s media (Sigma) supplemented with 10% heat-inactivated FBS (Gibco/Sigma) and 1X Pen/Strep (Gibco). 75μl of S2 media was used per larva with multiple larvae pooled per experiment. Cells in suspension were then transferred to glass-bottomed 96-well plates (Porvair) and allowed to adhere in a humidified box in the dark for 2 hours ahead of stimulation with heat-killed *S. cerevisiae* particles stained using calcofluor staining solution (Sigma).

*S. cerevisiae* (strain BY4741/accession number Y00000, Euroscarf consortium) were grown to exponential phase in YPD broth (Fisher) at 28°C. Yeast were heat killed at 60°C for 30 minutes, spun down and frozen at 20 × 10^9^ cells/ml. 1×10^9^ heat-killed yeast particles in 1ml of PBS (Fluka) were stained for 30 minutes at room temperature (with rotation) using 15μl of calcofluor staining solution. Stained yeast particles were washed in PBS and 1 × 10^6^ particles resuspended in 75μl S2 cell medium, which was then added to each well of larval hemocytes for 2 hours. Cells were fixed in wells using 4% EM-grade formaldehyde in PBS for 15 minutes and washed in PBS. Images were taken on a Nikon Ti-E inverted fluorescence microscope using a 20x objective lens and GFP and DAPI filter sets.

### Image analysis and statistical analysis

All microscopy images were processed using Fiji [92]. Images were typically analysed as maximum z-projections, with the exception of analysis of numbers of cells labelled via *VT-GAL4* lines (Figure 2h), wound responses (Figure 6c-d), vacuolation (Figure 6k) and quantification of ROS staining (Supplementary Figure 3f). Quantification was performed on blinded z-stacks for these analyses. Statistical tests were performed using Prism 7 (GraphPad Software, La Jolla, California, USA). P values less than 0.05 were deemed significant. A Student’s t-test was performed when comparing two sets of parametric data. When multiple comparisons were required, a one-way ANOVA with Dunnett’s multiple comparisons test was performed.

## Acknowledgements

This work was funded by a Wellcome/Royal Society Sir Henry Dale Fellowship (102503/Z/13/Z) awarded to IRE and a Bateson Centre studentship awarded to IRE, MPZ and JAC. Imaging work was performed at the Wolfson Light Microscopy Facility, using the Perkin Elmer spinning disk (MRC grant G0700091 and Wellcome grant 077544/Z/05/Z), Nikon A1 confocal/TIRF (Wellcome grant WT093134AIA) and Zeiss AiryScan microscopes. This work would not be possible without reagents and resources obtained from or maintained by the Bloomington *Drosophila* Stock Centre (NIH P40OD018537), the Vienna *Drosophila* Research Centre and Flybase (NIH and MRC grants U41 HG000739 and MR/N030117/1, respectively). We thank the *Drosophila* community for sharing *Drosophila* reagents (see Supplementary Table 1). We are grateful to Darren Robinson and the Fly Facility staff (University of Sheffield) for their support and to Phil Elks and Simon Johnston (University of Sheffield) for critical reading and feedback on the manuscript. We particularly thank Phil Elks and the now sadly departed Jarema Malicki for their advice and suggestions throughout the project. We thank Agata Grettka and Eleanor Castle for additional experiments that have contributed to our understanding of this project.

## Author contributions

Experiments performed by JAC, AB, ELA and IRE. JAC, AB and IRE wrote the initial manuscript. All authors contributed to experimental design and revision of the manuscript. The project was conceived and funding obtained by IRE and MZ.

## Declaration of interests

The authors declare no competing interests.

## Notes

### Competing Interest Statement

The authors have declared no competing interest.

