## Supplementary Information for "Identification of functionally-distinct macrophage subpopulations regulated by efferocytosis in *Drosophila*"

### Supplementary Methods

#### ROS staining of embryos

Embryos containing tdTomato-labelled plasmatocytes (with expression driven using *srp-GAL4* or *VT-GAL4*) were left in water for 30 minutes after dechorionation. Stage 15 embryos were then selected and transferred to a glass vial wrapped in foil containing 1ml peroxide-free heptane (supplier) and 1ml of 50μM dihydrorhodamine 123 (DHR123, Sigma-Aldrich) in PBS. Embryos were shaken at 250 rpm for 30 minutes. Following this, embryos were removed from the interface and mixed with halocarbon oil 700 (Sigma-Aldrich). Embryos were orientated individually in a droplet of this oil on a glass slide and then immediately imaged using a Zeiss Airyscan microscope (40x Plan-Apochromat oil immersion objective, NA=1.4), with z-spacing of 1μm and stacks totalling 30μm from the surface of the vitelline membrane down through the ventral nerve cord. Embryos were exposed to 10mM of H_2_O_2_ (Sigma-Aldrich) in PBS and peroxide-free heptane for 30 minutes prior to staining with DHR123 as a positive control. Negative control embryos were incubated in heptane (Sigma-Aldrich)/PBS (Oxoid) alone.

To quantify ROS levels the intensity of DHR123 staining was measured in the z-slice in which each macrophage exhibited its maximal cross-sectional area. The body of the macrophage was drawn around using the polygon tool in Fiji and then the area and mean gray value were measured in the GFP (DHR123) channel. Average mean gray value per plasmatocyte, per embryo was then plotted in Prism.

#### Morphological analysis of plasmatocytes

The vitelline membrane was manually removed from individual z-slices by drawing around the inside edge of the membrane with the freehand selection tool and using the clear outside command. Maximum projections were then created of the ventral midline region in Fiji. Following this, a region of interest was manually drawn around the area of individual plasmatocytes using the polygon tool and a range of cell shape descriptors and measurements calculated using Fiji.

#### Phagocytosis of *E. coli*

Dechorionated stage 15 embryos were mounted ventral-side up on a slide using double-sided Scotch tape, then dehydrated by incubating in a small container with silica beads for 7-8 minutes. Further dehydration was then prevented by covering embryos in a small drop of Voltalef oil. 1mg/ml pHrodo green *E. coli* BioParticles (Invitrogen; resuspended in PBS) were microinjected into the anterior of stage 15 embryos to determine the phagocytic capability of labelled plasmatocytes. Needles were created by pulling 15cm long 1mm glass capillaries (World Precision Instruments) using a Flaming/Brown P-1000 micropipette puller (Sutter, program 51). Needle tips were snapped using forceps under high magnification to create a bevelled end. Imaging was performed 1 hour after injection using an UltraView Perkin Elmer Spinning Disk system (40x UplanSApo oil immersion objective lens, NA 1.3). The proportion of *VT-GAL4* or *srp-GAL4* positive cells containing *E. coli* was scored.

### Supplementary Figure Legends

#### Supplementary Figure 1. Split GAL4 approach to label plasmatocyte subpopulations

(a) *srp-AD* and *VT-DBD* transgenic flies were generated: *srp-AD* drives expression of the activation domain (AD) of *GAL4* in all hemocytes (green), while VT enhancers drive expression of the DNA binding domain (DBD) of *GAL4* in a subpopulation of plasmatocytes and, in the case of some VT enhancers, other cell types (dark blue). (b-c) intersection of these expression patterns leads to the presence of both halves of *GAL4* (light blue) solely within those cells in which both *srp* and VT enhancer regions are active (c). This approach restricts activation of transgenes under *UAS* control (e.g. *UAS-stinger*) to those cells that are double positive for both *srp* and VT enhancer expression, ensuring expression is more specific to plasmatocyte subpopulations.

#### Supplementary Figure 2. *VT-GAL4* labelled subpopulations show no gross differences in morphology compared to non-labelled plasmatocytes

(a-e) representative images of plasmatocytes at stage 15 on the ventral midline labelled using the pan-hemocyte marker *srp-GMA* (green) and *UAS-tdTomato* (magenta) via *srp-GAL4* (a), *VT17559-GAL4* (b), *VT32897-GAL4* (c), *VT57089-GAL4* (d) and *VT62766-GAL4* (e); scale bars represent 20μm. Scatterplots showing plasmatocyte spread area (f), circularity (g), aspect ratio (h) and roundness (i) for positive control cells (labelled via *srp-GAL4*), and *VT-GAL4* positive/negative cells within embryos corresponding to those shown in (a-e). Cells positive for *VT-GAL4* driven tdTomato expression were compared to non-tdTomato expressing cells via Student’s t-test. No significant difference (ns) was found for any *VT-GAL4* line tested; lines and error bars represent mean and standard deviation, respectively; data points represent individual plasmatocytes taken from a minimum of 3 embryos. See Supplementary Table 1 for full list of genotypes.

#### Supplementary Figure 3. *VT-GAL4* labelled plasmatocytes show no gross differences in their ROS levels compared to the overall population

(a-e) representative images of *VT-GAL4* positive plasmatocytes (labelled via expression from *UAS-tdTomato*, magenta in merge) at stage 15 on the ventral midline stained via dihydrorhodamine 123 (DHR123) to show ROS levels (green in merge); *srp-GAL4* was used as a positive control to show the overall population. (f) scatterplot showing quantification of mean gray value per *srp-GAL4* or *VT-GAL4* labelled plasmatocyte, per embryo; lines and error bars represent mean and standard deviation, respectively. No statistically significant differences (p>0.05) were found between the *VT-GAL4* lines shown and the overall population (labelled via *srp-GAL4*) using a one-way ANOVA compared to control; n=12 (control), 11 (*VT17559-GAL4* and *VT32897-GAL4*), 14 (*VT57089-GAL4* and *VT62766-GAL4*) embryos; ns denotes not significantly different to control; scale bars represent 20μm (a-e). See Supplementary Table 1 for full list of genotypes.

#### Supplementary Figure 4. *VT-GAL4* labelled plasmatocytes show no gross differences in their phagocytosis of *E. coli* compared to the overall population

(a) ventral views of stage 15 embryos containing *srp-GAL4* (positive control, overall population) and *VT-GAL4* positive plasmatocytes (labelled via expression from *UAS-tdTomato*, magenta) 1 hour following injection with pHrhodo-labelled *E. coli* particles (green); *srp-GAL4* was used as a positive control to show the overall population; *srp-GMA* used to label *VT-GAL4* negative plasmatocytes. (b) shows zooms of *E. coli* positive plasmatocytes indicated by white boxes in (a). (c) scatterplot showing quantification of the proportion of pHrhodo *E. coli* positive plasmatocytes per embryo in the populations labelled via *srp-GAL4* or the indicated *VT-GAL4* reporter. Lines and error bars represent mean and standard deviation, respectively. No statistically significant difference (ns; p>0.05) was found between the *VT-GAL4* lines shown and plasmatocytes labelled via *srp-GAL4* using a Kruskal-Wallis test with a Dunn’s multiple comparison test (n=22, 31, 28, 29, 28); scale bars represent 20μm (a). See Supplementary Table 1 for full list of genotypes.

### Supplementary Movie Legends

#### Supplementary Movie 1. Plasmatocytes in similar positions within the embryo do not respond equally to inflammatory stimuli

GFP (green) and red stinger (magenta) labelled plasmatocytes responding to an epithelial wound at stage 15. Tracks of cell movements are shown via dots and lines. Magenta circles show cells responding to the wound; blue circles indicate cells that are the same distance from the wound, but fail to respond to the wound. Movie corresponds to stills shown in Figure 1a and lasts for 60 minutes post-wounding. Scale bar represents 20μm.

#### Supplementary Movie 2. Flow of *srp*-positive cells in circulation within a white pre-pupa

Movie showing movements of *srp*-positive cells within the hemolymph of a white pre-pupa. Cells labelled via *UAS-stinger* expression using the split GAL4 system in a positive control white pre-pupa (*w;srp-AD/UAS-stinger;srp-DBD/+*).

#### Supplementary Movie 3. Movement of *VT57089* subpopulation cells within a white pre-pupa

Movie showing movements of *VT57089* cells within the hemolymph of a white pre-pupa. Cells labelled via *UAS-stinger* expression using the split GAL4 system (*w;srp-AD/UAS-stinger;VT57089-DBD/+*). Movie plays twice with an overlay of the tracks of cells in circulation shown in repeat.

#### Supplementary Movie 4. Movement of *VT32897*-labelled, dorsal vessel-associated, non-hemocyte cells within a white pre-pupa

Movie showing rhythmic movements of *VT32897*-positive cells (likely to be nephrocytes) moving in time with pumping of the dorsal vessel in a white pre-pupa. Cells labelled via *UAS-stinger* expression using the split GAL4 system (*w;srp-AD/UAS-stinger;VT32897-DBD/+*).

### Supplementary Tables

#### Supplementary Table 1. Genotypes and sources of fly lines used in this study

#### Supplementary Table 2. Fly food recipe used in this study
