## Supplementary figures and images for "Identification of functionally-distinct macrophage subpopulations regulated by efferocytosis in *Drosophila*"

### Supplementary Figure 1

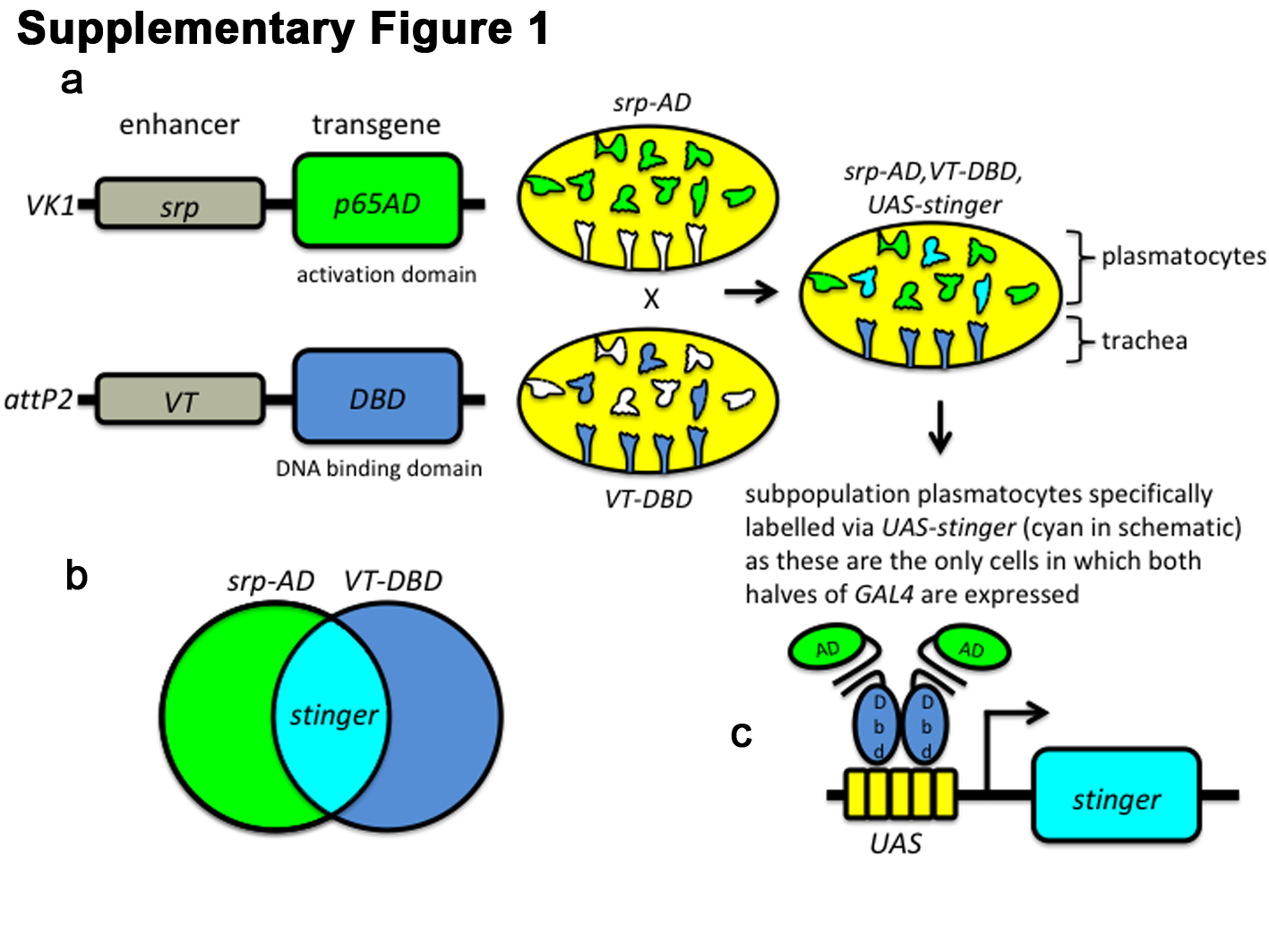

### Supplementary Figure 2

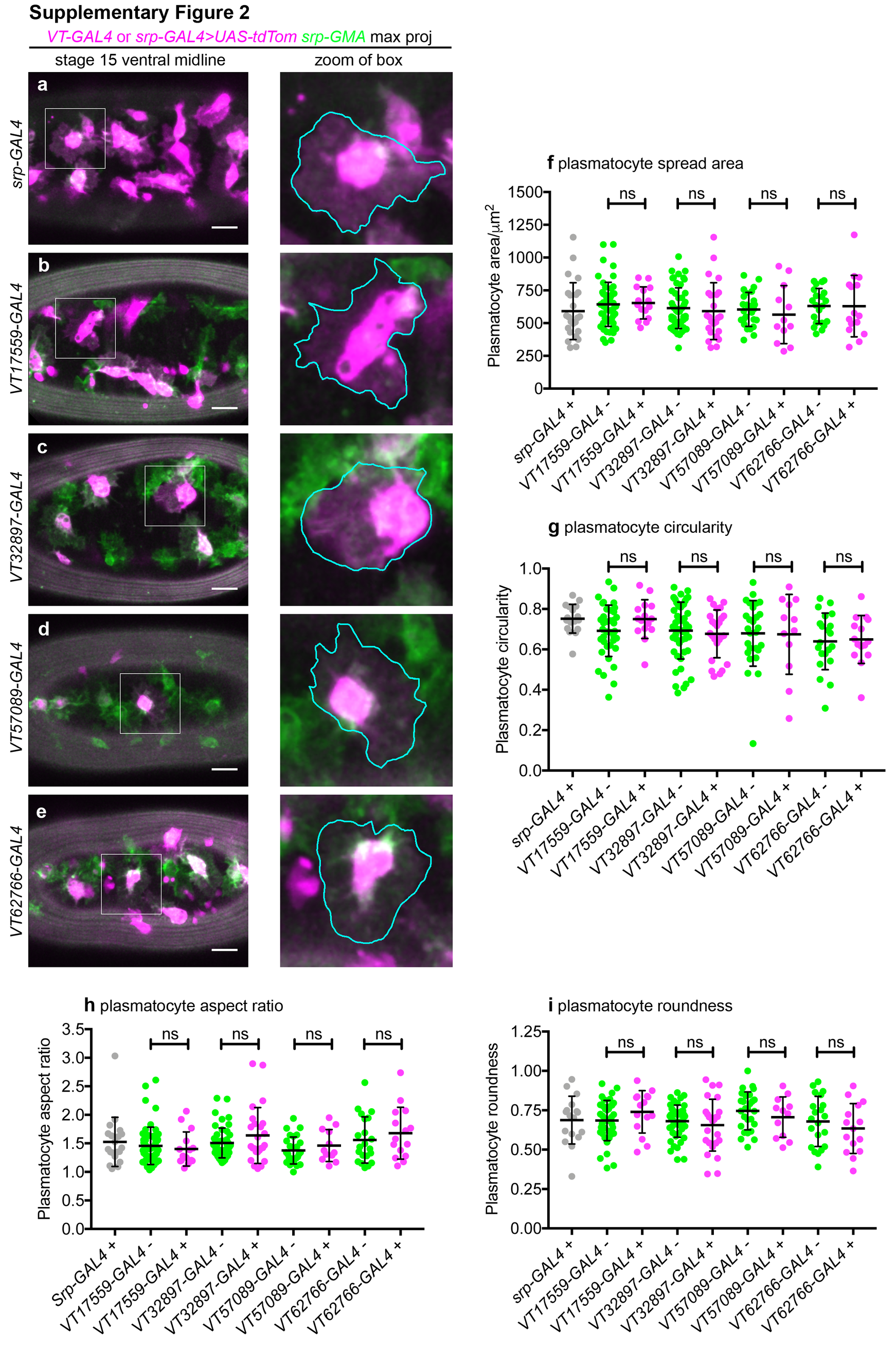

### Supplementary Figure 3

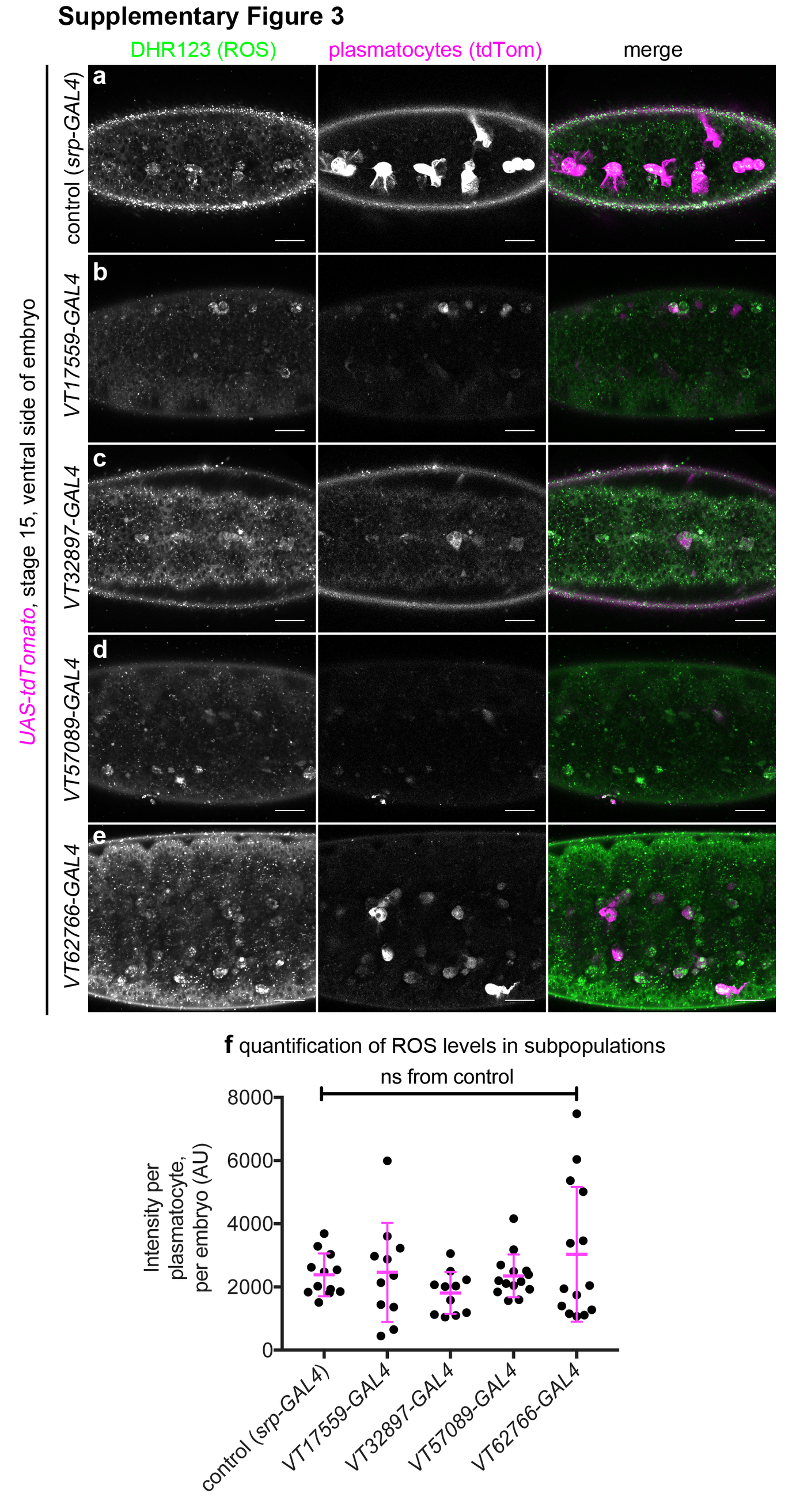

### Supplementary Figure 4

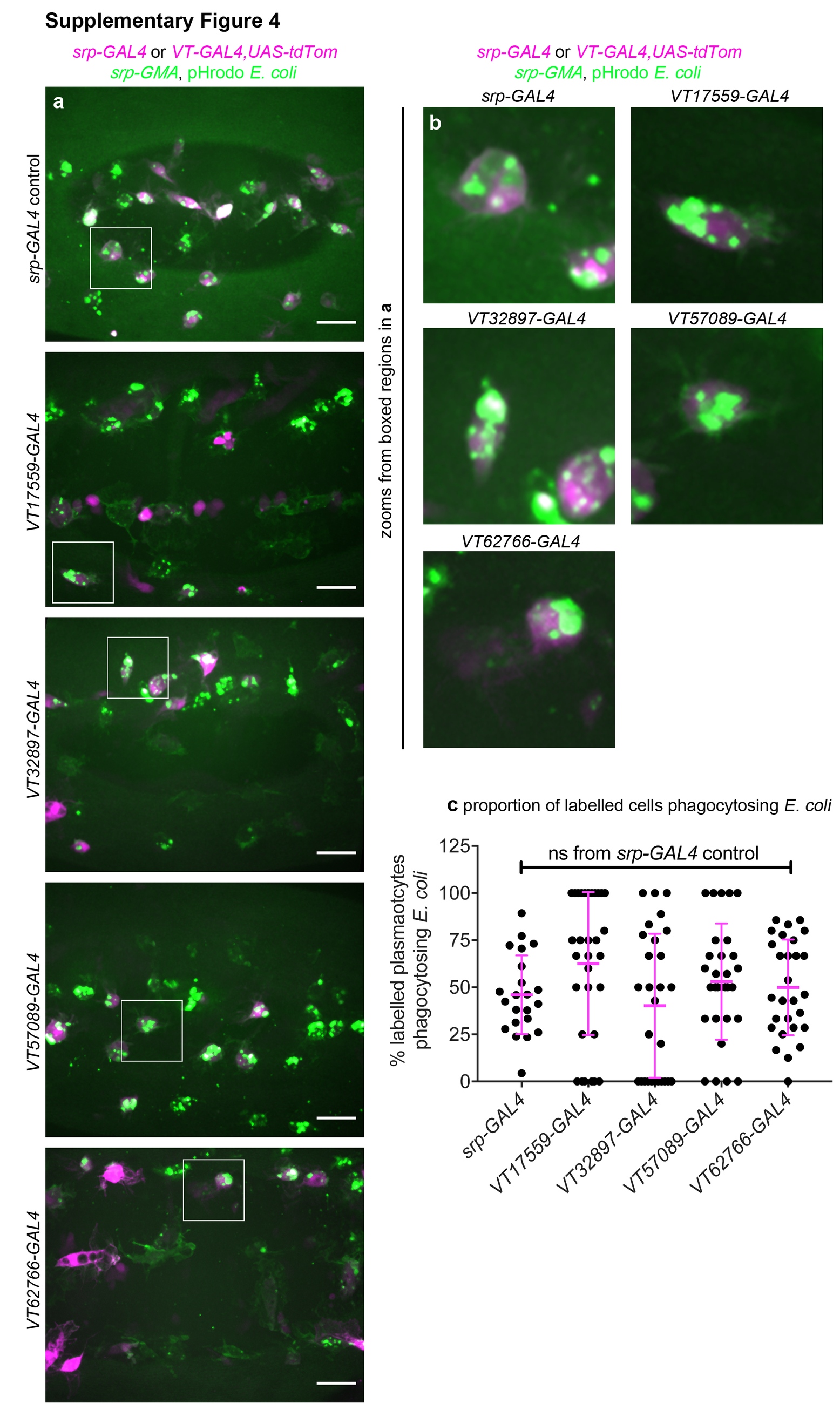
