## Supplementary Table 1 for "Identification of functionally-distinct macrophage subpopulations regulated by efferocytosis in *Drosophila*"

### Supplementary Table 1. Genotypes and sources of fly lines used in this study

| Figure | abbreviation | genotype | source | reference | notes |
|---|---|---|---|---|---|
| Figure 1a | *srp>GFP/srp>red stinger* | *w;srp-GAL4,UAS-GFP/srp-GAL4,UAS-red stinger* | from Will Wood, University of Edinburgh, UK | Bruckner et al., 2004 (*srp-GAL4*); Stramer et al., 2010 (*UAS-red stinger*) | |
| Figure 1b-d | *crq>GFP* | *w;crq-GAL4,UAS-GFP* | from Will Wood, University of Edinburgh, UK | Stramer et al., 2005 (*crq-GAL4*) | |
| Figure 1d | *srp-3xmCh/da>GC3ai* | *w;;srp-3x-mCherry/da-GAL4,UAS-GC3ai* | *srp-3x-mCherry* on chromosome 3 from Daria Siekhaus, IST Vienna, Austria; *UAS-GC3ai* from Magali Suzanne/Bruno Monier, Toulouse, France; *da-GAL4* from Alex Whitworth, University of Cambridge, UK | Gyoergy et al., 2018 (*srp-3x-mCherry*); Schott et al., 2018 (*UAS-GC3ai*); Wodarz et al., 1996 (*da-GAL4*) | |
| Figure 1e | *hml>GFP* | *w;hml(Δ)-GAL4,UAS-GFP;hml(Δ)-GAL4,UAS-GFP* | Bloomington Stock Centre (BL30140 and BL30142), Indiana, USA | Sinenko and Mathey-Prevot, 2004 | |
| Figure 2a | *VT41692-GAL4* | y w;;VT41692-GAL4 | Originally from VDRC, no longer available (available on request from I. Evans, University of Sheffield, UK) | Kvon et al., 2014 | in attP2 |
| Figure 2a | *VT17559-GAL4* | y w;;VT17559-GAL4 | Originally from VDRC, no longer available (available on request from I. Evans, University of Sheffield, UK) | Kvon et al., 2014 | in attP2 |
| Figure 2a | *VT32897-GAL4* | y w;;VT32897-GAL4 | Originally from VDRC, no longer available (available on request from I. Evans, University of Sheffield, UK) | Kvon et al., 2014 | in attP2 |
| Figure 2a | *VT57089-GAL4* | y w;;VT57089-GAL4 | Originally from VDRC, no longer available (available on request from I. Evans, University of Sheffield, UK) | Kvon et al., 2014 | in attP2 |
| Figure 2a | *VT62766-GAL4* | y w;;VT62766-GAL4 | Originally from VDRC, no longer available (available on request from I. Evans, University of Sheffield, UK) | Kvon et al., 2014 | in attP2 |
| Figure 2c | negative control (no *GAL4*) | *w;+/UAS-tdTomato;srp-GMA* | *UAS-tdTomato* from Bloomington Stock Centre (BL36327), Indiana, USA; *srp-GMA* on chromosome 3 from James Bloor, University of Kent, UK | Dutta et al., 2002 (*srp-GMA*) | |
| Figure 2c | positive control (*srp-GAL4*) | *w;srp-GAL4/UAS-tdTomato;srp-GMA* | | | |
| Figure 2e | *VT17559-GAL4* | *w;srp-GMA/UAS-tdTomato;VT17559-GAL4/srp-GMA* | *srp-GMA* on chromosome 2 from James Bloor, University of Kent, UK | Dutta et al., 2002 | |
| Figure 2e | *VT32897-GAL4* | *w;srp-GMA/UAS-tdTomato;VT32897-GAL4/srp-GMA* | | | |
| Figure 2e | *VT57089-GAL4* | *w;srp-GMA/UAS-tdTomato;VT57089-GAL4/srp-GMA* | | | |
| Figure 2e | *VT62766-GAL4* | *w;srp-GMA/UAS-tdTomato;VT62766-GAL4/srp-GMA* | | | |
| Figure 2f,h | *VT17559-GAL4* | *w;;VT17559-GAL4/UAS-GFP* | | | |
| Figure 2f,h | *VT32897-GAL4* | *w;;VT32897-GAL4/UAS-GFP* | | | |
| Figure 2f,h | *VT57089-GAL4* | *w;;VT57089-GAL4/UAS-GFP* | | | |
| Figure 2f,h | *VT62766-GAL4* | *w;;VT62766-GAL4/UAS-GFP* | | | |
| Figure 2g,h | *srp-GAL4* | *w;srp-GAL4/+;UAS-GFP/+* | | | |
| Figure 3-5 | $w^{1118}$ negative control | *w;UAS-stinger/+* | *UAS-stinger* from Bloomington Stock Centre (BL84277), Indiana, USA | | |
| Figure 3-5 | *srp-AD;srp-DBD* positive control | *w;srp-AD/UAS-stinger;srp-DBD/+* | made in this study (available on request from I. Evans, University of Sheffield, UK) | | *srp-AD* in VK1, *srp-DBD* in attP2 |
| Figure 3-5 | *srp-AD;VT17559-DBD* | *w;srp-AD/UAS-stinger;VT17559-DBD/+* | made in this study (available on request from I. Evans, University of Sheffield, UK) | | *srp-AD* in VK1, *VT17559-DBD* in attP2 |
| Figure 3-5 | *srp-AD;VT32897-DBD* | *w;srp-AD/UAS-stinger;VT32897-DBD/+* | made in this study (available on request from I. Evans, University of Sheffield, UK) | | *srp-AD* in VK1, *VT32897-DBD* in attP2 |
| Figure 3-5 | *srp-AD;VT57089-DBD* | *w;srp-AD/UAS-stinger;VT57089-DBD/+* | made in this study (available on request from I. Evans, University of Sheffield, UK) | | *srp-AD* in VK1, *VT57089-DBD* in attP2 |
| Figure 3-5 | *srp-AD;VT62766-DBD* | *w;srp-AD/UAS-stinger;VT62766-DBD/+* | made in this study (available on request from I. Evans, University of Sheffield, UK) | | *srp-AD* in VK1, *VT62766-DBD* in attP2 |
| Figure 5a | +ve con srp-3x-mCherry | *w;;+/srp-3x-mCherry* | *srp-3x-mCherry* on chromosome 2 from Daria Siekhaus, IST Vienna, Austria | Gyoergy et al., 2018 | |
| Figure 6 | *srp-GAL4* | *w;srp-GAL4/UAS-stinger* | | | |
| Figure 6 | *VT17559-GAL4* | *w;+/UAS-stinger;VT17559-GAL4/+* | | | |
| Figure 6 | *VT32897-GAL4* | *w;+/UAS-stinger;VT32897-GAL4/+* | | | |
| Figure 6 | *VT57089-GAL4* | *w;+/UAS-stinger;VT57089-GAL4/+* | | | |
| Figure 6 | *VT62766-GAL4* | *w;+/UAS-stinger;VT62766-GAL4/+* | | | |
| Figure 7b,c | control | *w;srp-GAL4,UAS-GFP/+;crq-GAL4,UAS-GFP/+* | | | |
| Figure 7b,c | *hc>cnx14D* | *w,UAS-cnx14D/w;srp-GAL4,UAS-GFP/+;crq-GAL4,UAS-GFP/+* | *UAS-cnx14D* from Harvard stock centre (d04188), Boston, USA | | |
| Figure 8a | control | *w;;crq-GAL4,UAS-GFP* | | | |
| Figure 8b | *repo* mutant | *w;;repo*$^{03702}$*,crq-GAL4,UAS-GFP* | *repo*$^{03702}$ from Bloomington Stock Centre (BL11604), Indiana, USA | Halter et al., 1995 | *ry*$^{506}$ allele recombined off (Armitage et al., 2020 Bioarchive) |
| Figure 8c | negative control | *w;srp-H2A-3x-mCherry/+* | *srp-H2A-3x-mCherry* on chromosome 2 from Daria Siekhaus, IST Vienna, Austria | Gyoergy et al., 2018 | |
| Figure 8d,h | *VT17559-GAL4* control | *w;UAS-stinger/srp-H2A-3x-mCherry;VT17559-GAL4/+* | | | |
| Figure 8d',h | *VT17559-GAL4,repo* | *w;UAS-stinger/srp-H2A-3x-mCherry;VT17559-GAL4,repo*$^{03702}$*/repo*$^{03702}$ | | | |
| Figure 8e,h | *VT32897-GAL4* control | *w;UAS-stinger/srp-H2A-3x-mCherry;VT32897-GAL4/+* | | | |
| Figure 8e',h | *VT32897-GAL4,repo* | *w;UAS-stinger/srp-H2A-3x-mCherry;VT32897-GAL4,repo*$^{03702}$*/repo*$^{03702}$ | | | |
| Figure 8f,h | *VT57089-GAL4* control | *w;UAS-stinger/srp-H2A-3x-mCherry;VT57089-GAL4/+* | | | |
| Figure 8f',h | *VT57089-GAL4,repo* | *w;UAS-stinger/srp-H2A-3x-mCherry;VT57089-GAL4,repo*$^{03702}$*/repo*$^{03702}$ | | | |
| Figure 8g,h | *VT62766-GAL4* control | *w;UAS-stinger/srp-H2A-3x-mCherry;VT62766-GAL4/+* | | | |
| Figure 8g',h | *VT62766-GAL4,repo* | *w;UAS-stinger/srp-H2A-3x-mCherry;VT62766-GAL4,repo*$^{03702}$*/repo*$^{03702}$ | | | |

| Figure | abbreviation | genotype | source | reference | notes |
|---|---|---|---|---|---|
| Supplementary Figures 2-4 | *srp-GAL4*/control | *w;srp-GAL4/UAS-tdTomato;srp-GMA* | | | + refers to cells with tdTomato and GMA expression |
| Supplementary Figures 2-4 | *VT17559-GAL4* | *w;srp-GMA/UAS-tdTomato;srp-GMA/VT17559-GAL4* | | | + refers to cells with tdTomato and GMA expression, - refers to cells with GMA expression but without tdTomato |
| Supplementary Figures 2-4 | *VT32897-GAL4* | *w;srp-GMA/UAS-tdTomato;srp-GMA/VT32897-GAL4* | | | + refers to cells with tdTomato and GMA expression, - refers to cells with GMA expression but without tdTomato |
| Supplementary Figures 2-4 | *VT57089-GAL4* | *w;srp-GMA/UAS-tdTomato;srp-GMA/VT57089-GAL4* | | | + refers to cells with tdTomato and GMA expression, - refers to cells with GMA expression but without tdTomato |
| Supplementary Figures 2-4 | *VT62766-GAL4* | *w;srp-GMA/UAS-tdTomato;srp-GMA/VT62766-GAL4* | | | + refers to cells with tdTomato and GMA expression, - refers to cells with GMA expression but without tdTomato |
| Supplementary Movie 1 | *srp>GFP/srp>red stinger* | *w;srp-GAL4,UAS-GFP/srp-GAL4,UAS-red stinger* | | | |
| Supplementary Movie 2 | *srp* | *w;srp-AD/UAS-stinger;srp-DBD/+* | | | |
| Supplementary Movie 3 | *VT57089* | *w;srp-AD/UAS-stinger;VT57089-DBD/+* | | | |
| Supplementary Movie 4 | *VT32897* | *w;srp-AD/UAS-stinger;VT32897-DBD/+* | | | |

nb1 $w$=$w^{1118}$ allele unless stated
nb2 source/reference/notes only shown on first instance in table
