## Supplementary Table 2 for "Identification of functionally-distinct macrophage subpopulations regulated by efferocytosis in *Drosophila*"

### Supplementary Table 2. Fly food recipe used in this study

| Ingredient | Amount | Manufacturer | Supplier |
|---|---|---|---|
| Cold tap water | 1 Litre | | |
| Medium Cornmeal | 80g | Triple Lion | Lembas/Easton Enterprises, Sheffield, UK |
| Dried Yeast | 18g | Kerry Ingredients | BTP Drewitt, London, UK |
| Soya Flour | 10g | Lembas Wholefoods | Lembas |
| Malt Extract | 80g | Rayner’s Essentials | Lembas |
| Molasses | 40g | Rayner’s Essentials | Lembas |
| Agar | 8g | | BTP Drewitt |
| 10% Nipagin in Absolute Ethanol | 25ml | Clariant UK Ltd Fisher | Chemlink Specialities Ltd, Manchester, UK Fisher |
| Propionic Acid | 4ml | Fisher | Fisher |
